# Persistence of parental age effect on somatic mutation rates across generations in *Arabidopsis*

**DOI:** 10.1101/2022.01.12.476102

**Authors:** Shashi Bhushan, Amit Kumar Singh, Yogendra Thakur, Ramamurthy Baskar

## Abstract

In the model plant *Arabidopsis thaliana*, parental age is known to affect somatic mutation rates in their immediate progeny and here we show that this age dependent effect persists across successive generations. Using a set of detector lines carrying the mutated *uidA* gene, we examined if a particular parental age maintained across five consecutive generations affected the rates of base substitution (BSR), intrachromosomal recombination (ICR), frameshift mutation (FS), and transposition. The frequency of functional GUS reversions were assessed in seedlings as a function of identical/different parental ages across generations. In the context of a fixed parental age, BCR/ICR rates were unaffected in the first three generations, then dropped significantly in the 4^th^ and increased in most instances in the 5^th^ generation. On the other hand, with advancing parental ages, BSR/ICR rates remained high in the first two/three generations, with a striking resemblance in the pattern of mutation rates. We adopted a novel approach of identifying and tagging flowers pollinated on a particular day, thereby avoiding biases due to potential emasculation induced stress responses. Our results suggest a time component in counting the number of generations a plant has passed through self-fertilization at a particular age in determining the somatic mutation rates.

## Introduction

Adaption during evolution heavily relies on variations within the DNA sequence (Futuyma, 2013) and somatic mutations are an important source of genetic variation in plants (Otto and Gerstein, 2006; Eyre-Walker and Keightley, 2007). A fraction of such spontaneous mutations are known to be transmitted from reproductive tissues arising later in development (Bilichak and Kovalchuk, 2016) and thus, it is important to understand the frequency and regulators of such events. Spontaneous mutations largely arise as a consequence of impaired proof-reading activity and errors occurring during DNA repair (Golubov *et al.,* 2010; Martina *et al.,* 2012).

In *A. thaliana,* the single nucleotide mutation (SNM) rate is around 6.95 × 10^-9^ per site per generation, occurring mostly within transposable elements (TEs) residing in centromeric regions (Weng *et al.,* 2019).

Stress of different kind experienced by organisms are known to affect the phenotypic traits of the offspring. Such environmental influences are known to be transmitted and termed transgenerational plasticity or parental effects (Tariel *et al.,* 2020). In certain situations, plants subjected to a particular stress are known to produce offspring that are better adapted to the same stress (Sultan, 1996; Galloway and Etterson, 2007; Holeski, 2007; Salinas and Munch, 2012). Thus the phenotype of an organism is also determined by the impact of various environmental/growth conditions experienced in earlier generations (Agrawal *et al.,* 1999; Galloway and Etterson, 2007; Salinas *et al.,* 2013).

Plants are constantly exposed to biotic and abiotic stresses, triggering different classes of somatic mutations (Molinier *et al.,* 2006). Such experiences are recorded in the somatic cell lineage, affecting the genome stability to confer better adaptation to the same stress in subsequent generations (Molinier *et al.,* 2006; Boyko *et al.,* 2007; Boyko and Kovalchuk, 2010; Bilichak and Kovalchuk, 2016). Thus, any reproductive cell derived from such somatic tissues would also transfer this recorded memory to the subsequent generation (Molinier *et al.,* 2006). This phenomenon of the acquired memory is often referred to as transgenerational inheritance, and is defined as the ability of an organism to ‘remember’ the environmental conditions of the past at a molecular level, resulting in a subsequent change in the phenotype of the progeny (Tricker, 2015).

The induced somatic mutation rates in *Arabidopsis* are reported to persist for up to 4^th^ generations (Molinier *et al.,* 2006), but other studies contradict the existence of this phenomenon (Pecinka *et al.,* 2009; Groot *et al.,* 2016). Furthermore, there are also reports suggesting that such transgenerational effects depend on the nature of the stress (Pecinka *et al.,* 2009). Although transgenerational effects are known to be inherited through both the parents in a dominant manner (Molinier *et al.,* 2006), some studies indicate that the influence of maternal gametes on the transgenerational effect is more significant than paternal gametes (Mosher, 2010; Boyko and Kovalchuk, 2010). Transgenerational effects are context-dependent and their reproducibility is reported to be low (Groot *et al.,* 2016).

Previously, we reported that parental age of *Arabidopsis* influenced somatic mutation rates in their immediate progeny (Singh *et al.,* 2015) and here demonstrate the persistence of this effect in its transgenerational progenies as well. Parental age at the time of reproduction has a strong influence on the patterns of somatic mutation across generations and this age-dependent information persists for a few generations after which there seems to be resetting of mutation s rates.

## Results

Using a set of *Arabidopsis* mutation detector lines (Liu and Crawford, 1998; Li *et al.,* 2004; Azaiez *et al.,* 2006; Auwera *et al.,* 2008), we examined if a particular parental age affected the spontaneous base substitution rates (BSR), intrachromosomal recombination (ICR), frameshift mutation (FS) and transposition rates in the next five consecutive generations. Self-pollinated plants of four different ages (38, 43, 48, and 53 days after sowing (DAS)) were grown, following which the respective reproductive ages were retained in the subsequent generations also (Fig. 1). The mutation rates were examined in two different ways. 1. Counting reversion events in seedlings obtained from plants of the same age set as depicted in Fig. 1. For instance, mutation rates were compared in seedlings of 38 DAS plants in five consecutive generations (comparison of same age (same colors) shown in Fig. 1). 2. Furthermore, we compared the mutation rates as a function of advancing parental age across generations. For eg., the mutations rates were compared across the different ages (colors) of the same generation, i.e. 38, 43, 48, and 53 DAS plants among F1 as in Fig. 1.

**Figure 1.**
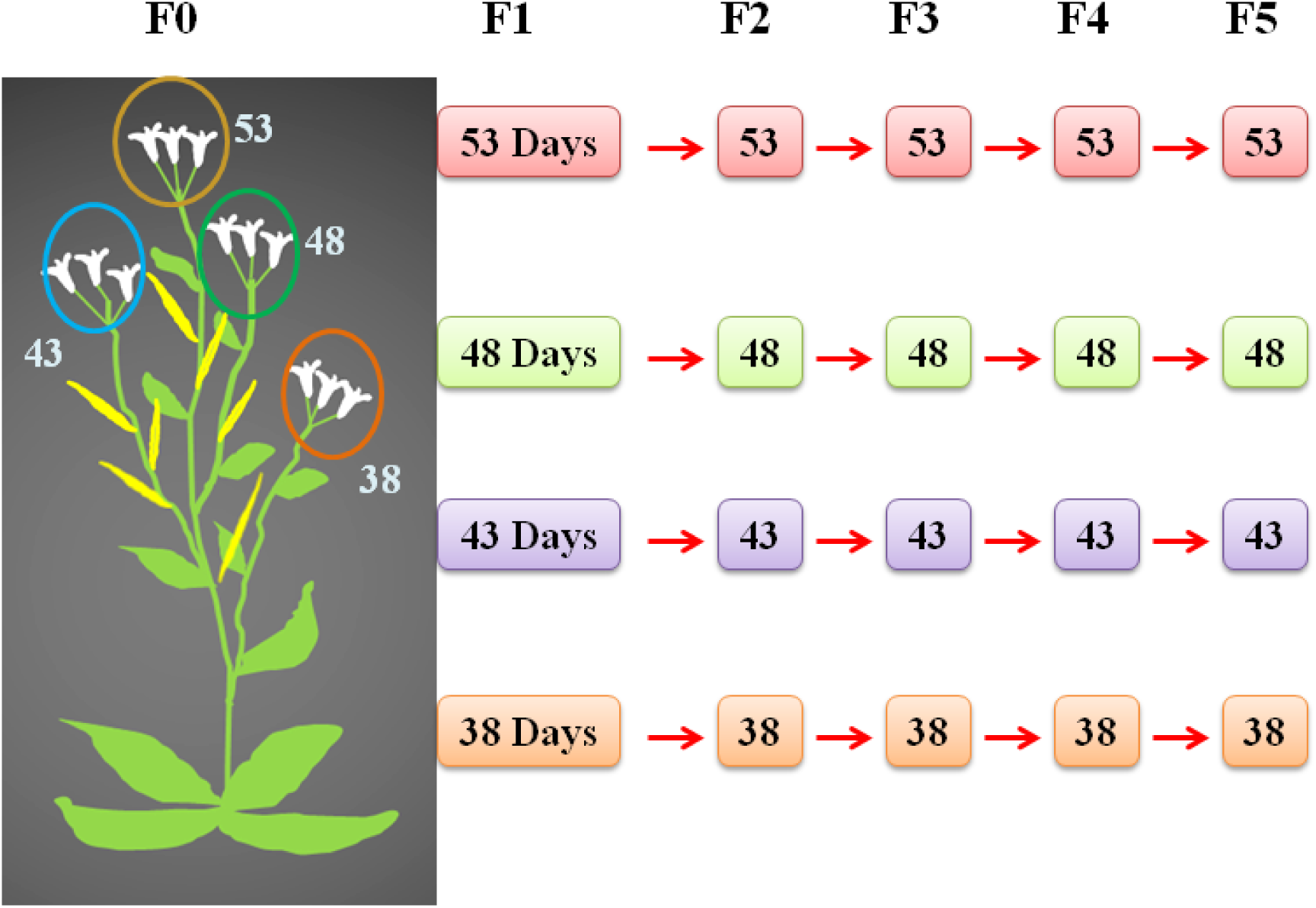
Selfing scheme: Flowers of different DAS were marked with different colored threads and in the subsequent generations also, the self-pollinating age was retained.

### Normalization of mutation rates after correcting the cell number and ploidy levels

Mutation rates were calculated based on the reversion of a mutated or truncated *uidA* gene to wild type sequence whereby acquisition of functional GUS activity results in the development of blue colored spots upon histochemical staining. Thus, the overall reversion frequency was estimated via quantification of the number of such blue spots with respect to the total number of seedlings screened. Since ploidy levels, cell size, and their number vary significantly from seedlings obtained from plants of different ages (Singh *et al.,*, 2015), normalization of genome per nucleus is required. Plant cells are known to be uneven in size even within a particular tissue (Sugimoto-Shirasu and Roberts, 2003). In the leaf epidermal cells of *Arabidopsis,* a strong correlation exists between the size of the cell and the amount of DNA (Melaragno *et al.,* 1993). Plant cells frequently undergo endoreduplication, resulting in higher ploidy per nucleus which is correlated with increased cell size as well (Sugimoto-Shirasu and Roberts, 2003; Singh *et al.,* 2019). As a result, the individual cell size and their numbers significantly contribute to the overall leaf size (Tsukaya, 2003). As the ploidy per nucleus, the cell size, and cell number can vary in seedlings derived from parents of different ages/generations, it is essential to normalize the number of genomes per nucleus. As the leaf size is largely determined by the epidermis (Savaldi-Goldstein *et al.,* 2007; Marcotrigiano, 2010), the fourth true leaf (excluding the cotyledons) of Columbia (Col-0) wild type plants were used to determine cell size and number (Singh *et al.,* 2015). We used a scanning electron microscope (SEM) to count the number of adaxial epidermal cells in a specified area of the leaf and measured the cell size. Thus, the mutation rates were normalized per haploid genome as estimated from cell number and average ploidy in the 4^th^ true leaf.

As a function of constant or advancing parental ages, there was wide variation in the epidermal cell number (Fig. 2, E, F), and such differences were not apparent in the context of cell size (Fig. 2, G, H). Interestingly, as a function of fixed parental age, the epidermal cell number increases in the 4^th^ and drops significantly in the 5^th^ generation (Fig. 2E), although this largely depends on the age group. Consistently across all parental ages, the fraction of 2X nuclei increased in the fourth generation alone (Fig. 3A) and the percentage of nuclei with different ploidy levels remained more or less constant in the first three age sets (Fig.3B) regardless of the generation. Depending on age/generation, there were differences in ploidy levels per leaf cell nucleus, and in older ages (53 DAS), the variations in ploidy levels were highest compared to susbequent generations (Fig. 3C). Hence, the mutation rates were corrected for differences in genome numbers (obtained from average ploidy per nucleus) and cell numbers (Table S1).

**Figure 2.**
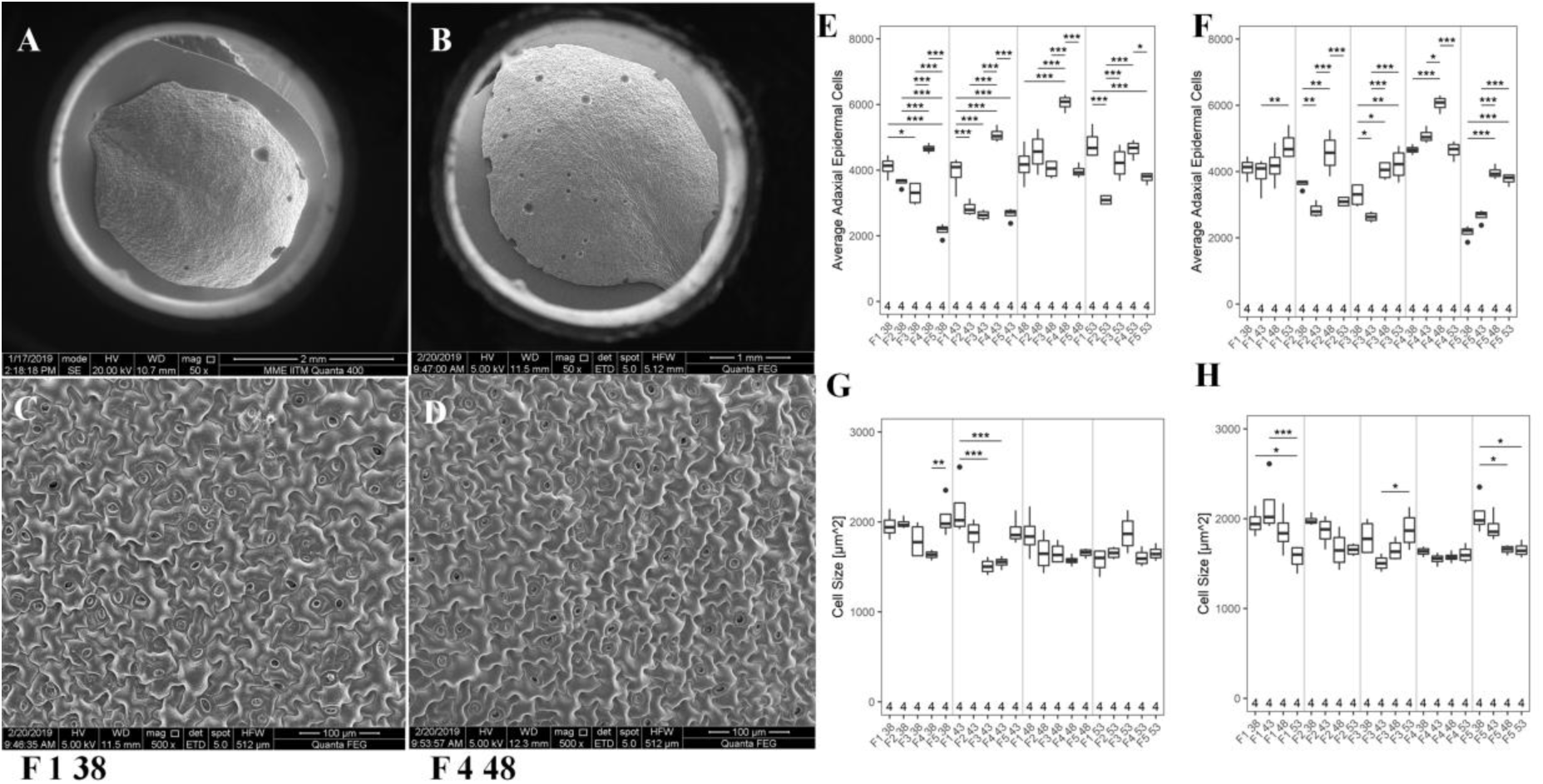
SEM image of the 4^th^ true leaf from seedlings of A. F1 (38 DAS) and B. F4 (48 DAS) generations. Differences in leaf size (A-B) and cell size (C-D) from seedlings of A and B. E-F: The average adaxial epidermal cell number. G-H: The average cell size in three-week-old seedlings derived from parents of (G) identical age sets and (H) different ages. The numbers in X-axis indicates the biological replicates analyzed. *P* values were corrected for multiple testing. No asterisk represents, no significant differences. *, *P <* 0.05; **, *P* < 0.01; ***, *P* < 0.001.

**Figure 3.**
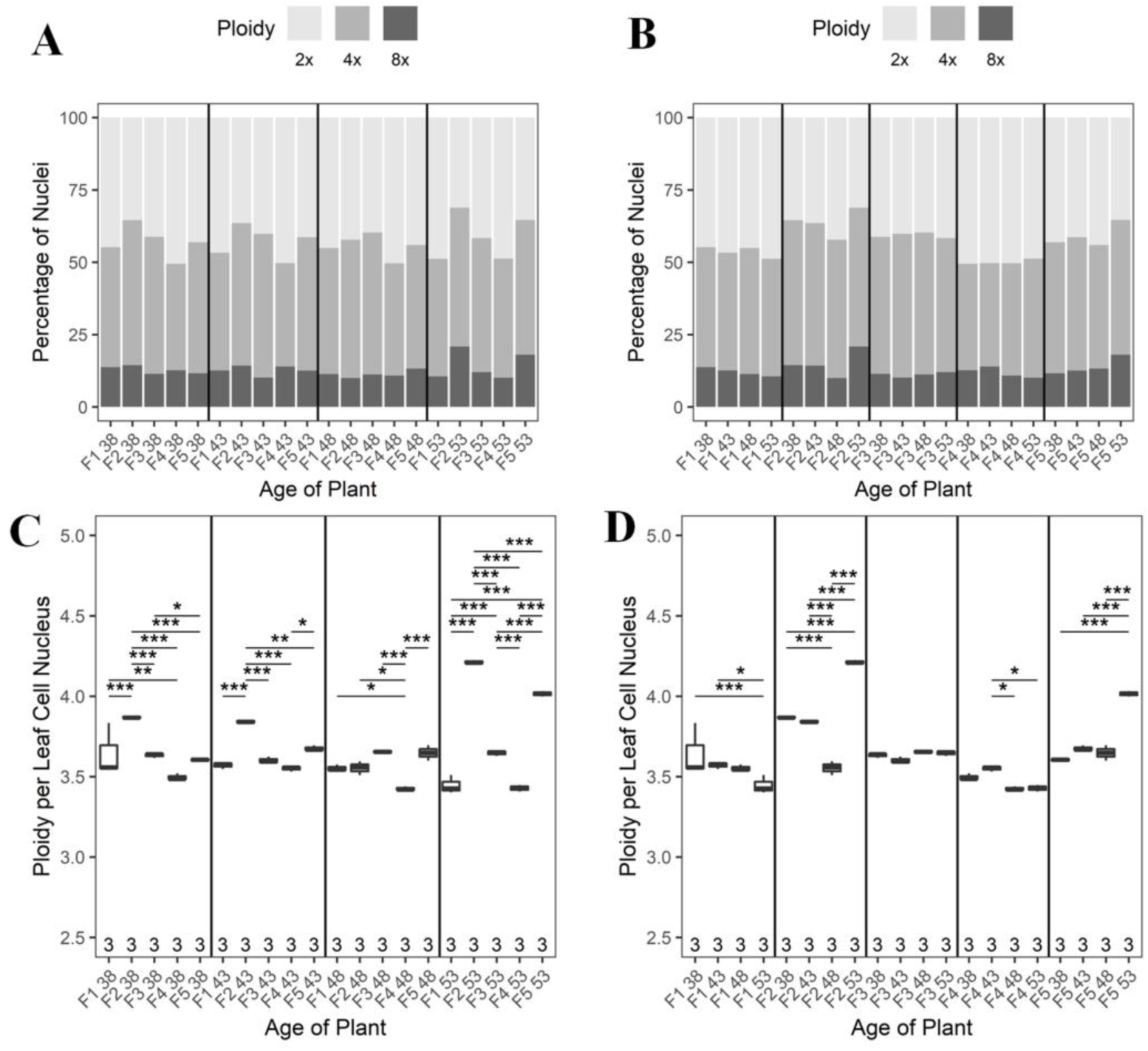
Comparison of ploidy levels across generations. The percentage of diploid, tetraploid, and octoploid nuclei as a function of (A) same and (B) advancing parental ages. Different shades of grey represent changes in ploidy levels. C-D: Ploidy per leaf cell nucleus as a function of C. Constant parental age, D. Different parental ages. The numbers along the X-axis of the graph shows the analyzed biological replicates. *P* values were corrected for multiple testing. No asterisk represents, no significant differences. *, *P <* 0.05; **, *P* < 0.01; ***, *P* < 0.001.

### Comparsion of reversion rates after manual and natural selfing

Throughout the study, self pollinated flowers of particular DAS were tagged with colored threads to avoid a potential emasculation indued stress response which may distort the mutation rates (Fig 4). Accordingly, the mutation rates were compared between two sets of seedlings one derived from individual plants manually self-pollinated on different DAS, while the other was obtained from a single plant, but from flowers of different DAS marked with colored threads. Although the pattern of mutation rates after normalization were largely similar, a 0.2 to 0.5 fold higher BSR, ICR, and FS mutation rates were observed in seedlings from manually pollinated plants of advanced ages (Fig 4) and this increased reversion rates may possibly be due to the result of repeated physical contacts followed by emasculation triggering a stress response.

**Figure 4.**
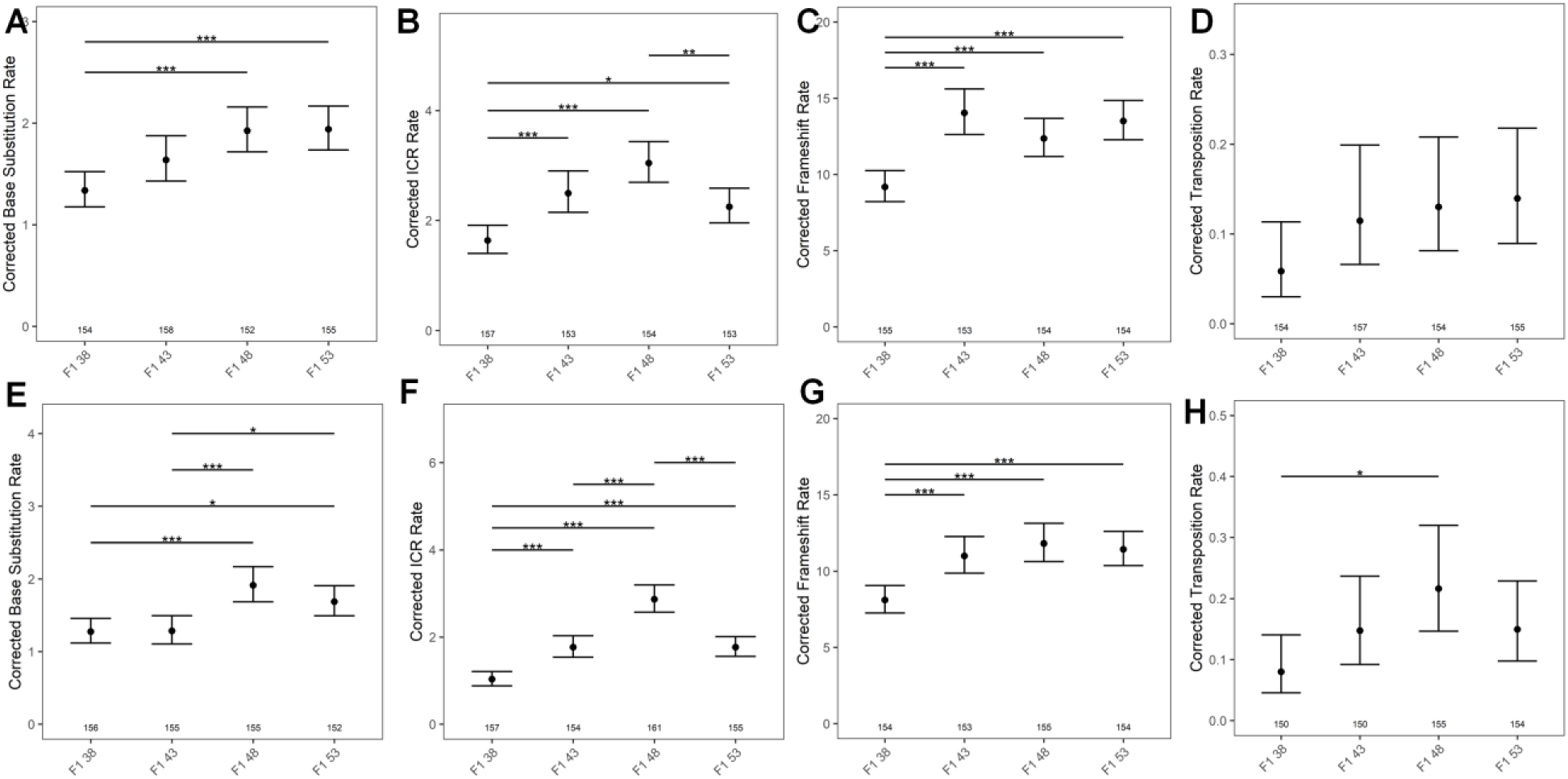
Two methods of selfing and its impact on mutation rates. The top panel represents somatic mutation rates after emasculation followed by manual selfing (A) Base Substitution Rates (BSR), (B) Intrachromosoal Recombination (ICR), (C) Frameshift (FS) and (D) Transposition and the panel below represents somatic mutation rates subsequent to marking self-fertilized flowers of particular DAS. (E) BSR, (F) ICR, (G) FS and (H) Transpotion. The numbers along the X axis shows the number of seedlings analyzed. *P* values were corrected for multiple testing. No asterisk represents, no significant differences. *, *P <* 0.05; **, *P* < 0.01; ***, *P* < 0.001.

### As a function of constant parental age, C to T transitions (BSR) are stable in first three consecutive generations

To determine the frequency of C to T transitions as a function of constant or different parental ages across generations, we used the 1390_T-C_ detector line. In the open reading frame of the *uidA* gene, T is mutated to C at the 1390^th^ position (Van der Auwera *et al.,* 2008). In seedlings of 38, 43, and 48 DAS plants, the variation in BSR rates were not significant in the first three generations. However, in the 4^th^ generation, seedlings from 38 and 48 DAS plants exhibited a significant drop in BSR, and thereafter, a significant increase in the 5^th^ generation (Fig. 5A). Similarly, in seedlings from 43 DAS plants, a surge in BSR was observed in the 5^th^ generation (Fig. 5A). Except for the seedlings obtained from the 53 DAS plants, the BSR increased in the 5^th^ generation consistently. Taken together, our results suggest that BSR significantly depends on the number of generations a plant has undergone selfing at a particular age.

**Figure 5.**
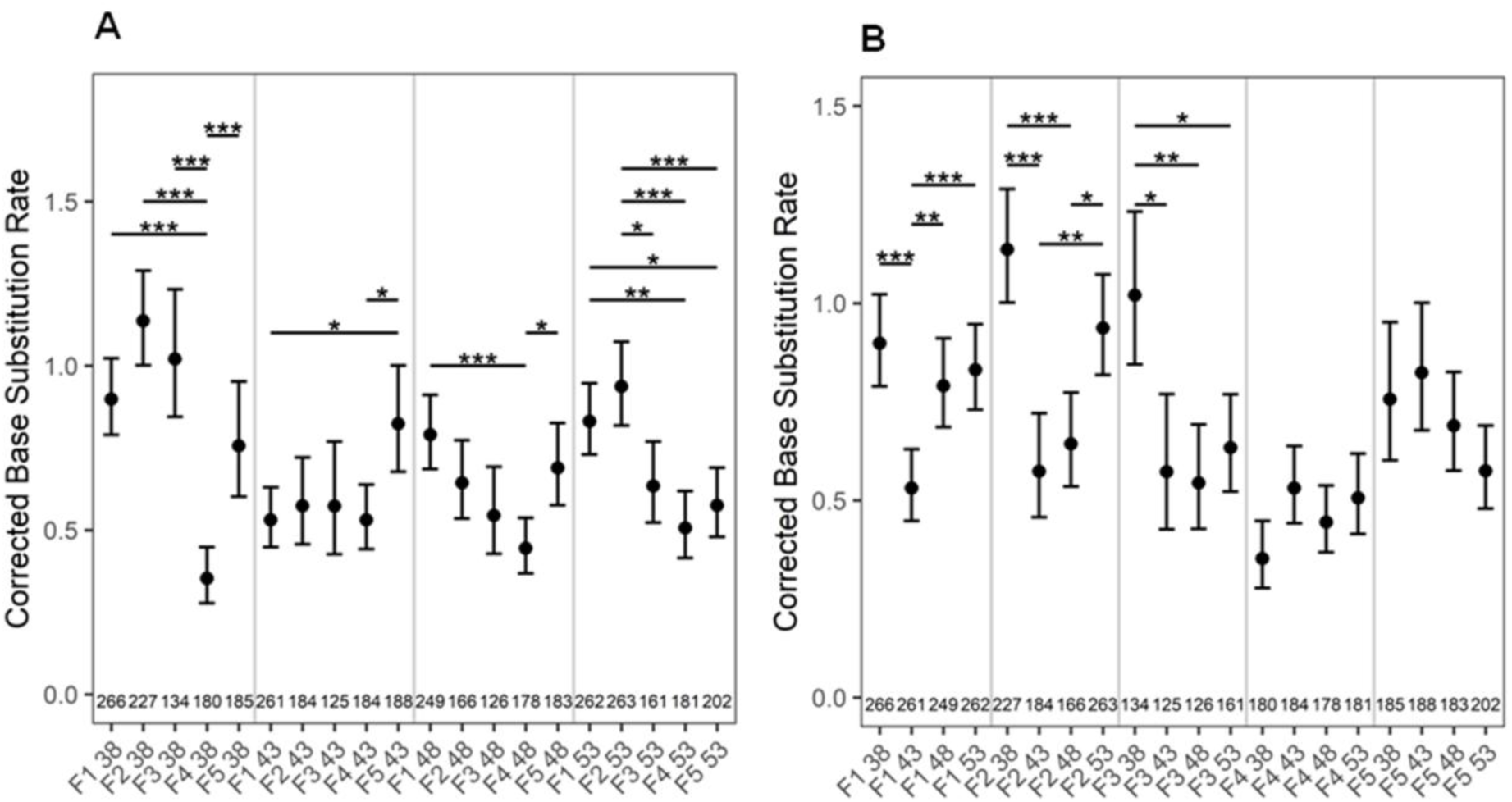
A-B: BSR as a function of A. constant, B. different parental age across different generations. The numbers along the X axis shows the number of seedlings analyzed. *P* values were corrected for multiple testing. No asterisk denotes no significant difference *. *P <* 0.05; **, *P* < 0.01; ***, *P* < 0.001.

### Younger parents produce offspings with more BSR

By comparing mutation rates across different ages in five consecutive generations, we observed high BSR in the first three generations of seedlings from 38 DAS plants compared to other ages (Fig. 5B). The BSR dropped significantly in seedlings of 43 DAS plants, and increased thereafter as a function of advancing parental ages (48 and 53 DAS). Surprisingly, this pattern was observed only in the first two generations. In the 4^th^ and 5^th^ generations, there was no significant change in BSR, irrespective of parental age (Fig. 5B). These results reiterate the notion that BSR is strongly impacted by the the number of generations a plant has gone through selfing at a particular age.

### ICR rates are stable as a function of constant parental age in three consecutive generations

To examine the effect of reproductive age on ICR rates across generations, we utilized the detector line R2L1, containing two inverted catalase introns within the gene *uidA*. In this system, a recombination event between similar sequences of the catalase intron would result in a functional *uidA* gene, leading to the GUS expression (Li *et al.,* 2004). With a constant age of 38 DAS, there was no significant change in ICR in the first four generations, but the rates significantly increased in the 5^th^ generation (Fig. 6A). However, when the parental age was retained at 43 or 48 DAS, there was no change in ICR rates in the first three generations, following which it decreased in 4^th^ but again surged in the 5^th^ generation. Although such a pattern was also observed in seedlings obtained from 53 DAS plants, the increase in ICR rates was not significant in the 5^th^ generation.

**Figure 6.**
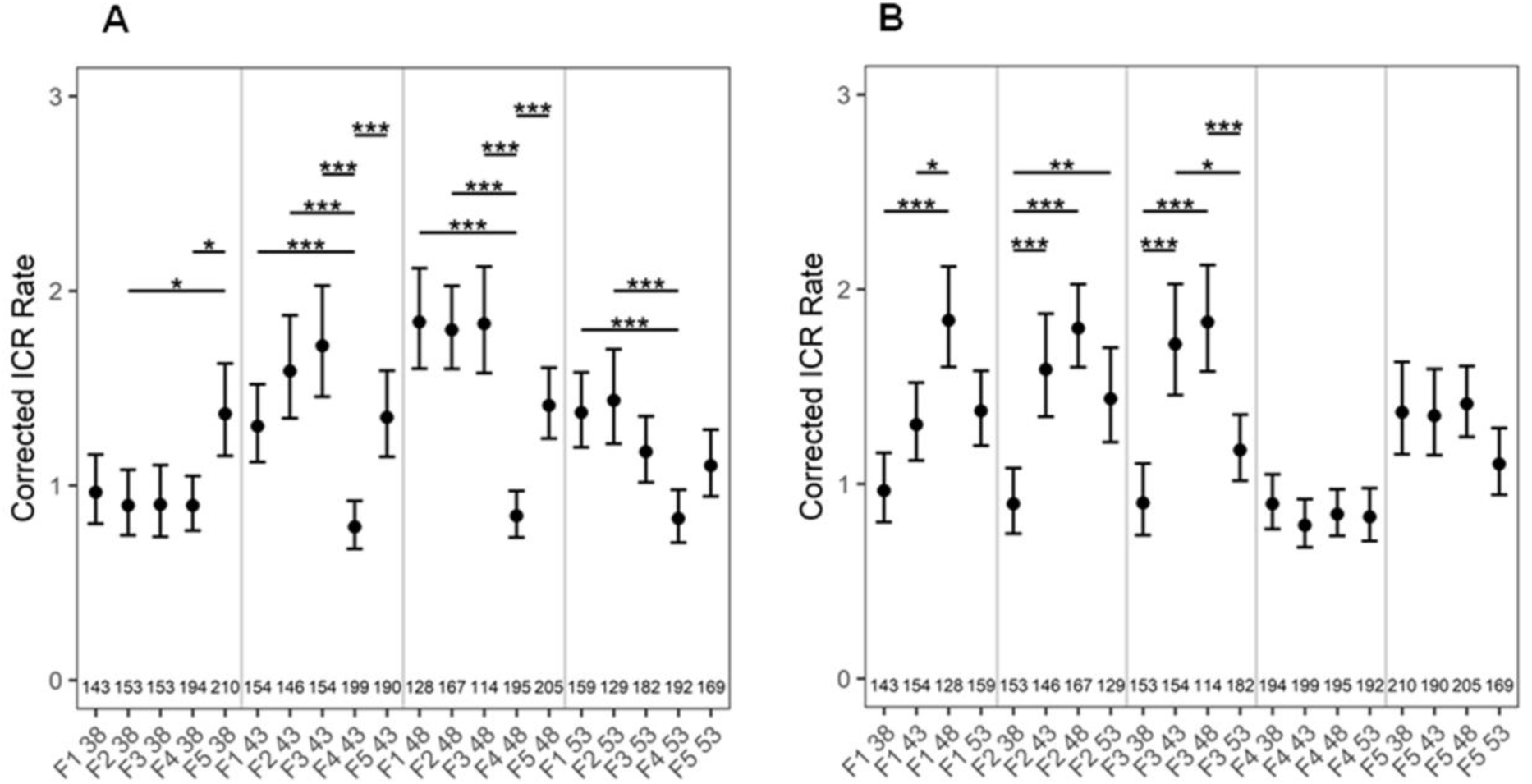
A-B: ICR rates as a function of A. constant, B. increasing parental age across different generations. The numbers at the bottom of the graph show the number of seedlings analyzed. *P* values were corrected for multiple testing. No asterisk represents no significant difference. *, *P <* 0.05; **, *P* < 0.01; ***, *P* < 0.001.

Strikingly, like BSR, ICR rates shoot up in the 5^th^ generation following the aforementioned pattern, and this phenomenon was observed across all ages. This suggests that ICR rates is determined by the number of generations a plant has been previously selfed at a particular reproductive age (Fig. 6A). Interestingly, there is a sigmoidal distribution of mutation rates upon comparison between different age sets across all generations (Fig. 6A).

### ICR rates increases with increasing parental age but drops in very old ages

Although, the increase in ICR rates (from seedlings of 38, 43, and 48 DAS) and the drop (in 53 DAS) were apparent in the first three generations, this pattern was not observed in the subsequent two generations (Fig. 6B). Thus, these observations suggest that the influence of advanced age on ICR rates is stronger in the first three generations, following which the influence gets weaker.

### Frameshift rates show a stochastic pattern as a function of constant parental age regardless of generation

To score FS mutations as a function of constant or advancing parental ages, transgenic Col-0 plants containing out-of-frame guanine repeats (G10) in the *uidA* reporter gene (Azaiez *et al.,* 2006) were used. The *uidA* gene function is restored either by the addition of two or deletion of one guanine nucleotide. In the seedlings of 38, 43, or 48 DAS plants, FS rates dropped significantly from the 3^rd^ generation and this trend persists across subsequent generations or undergoes further reduction in later generations. However, the FS rates always increased in the 5^th^ generation (Fig. 7A). An exception to this pattern was from the seedlings of 53 DAS plants, where FS rates dropped significantly from the 2^nd^ and lasted till the 5^th^ generation (Fig. 7A). Like BSR/ ICR, FS rates also shoots up significantly in the 5^th^ generation, suggesting a time component in counting the number of generations a plant has passed through self-fertilization. If FS rates were compared as a function of advancing parental age across generations, the rates remained almost constant (38/48 DAS), or gradually declined with advancing age (53 DAS) (Fig. 7B).

**Figure 7.**
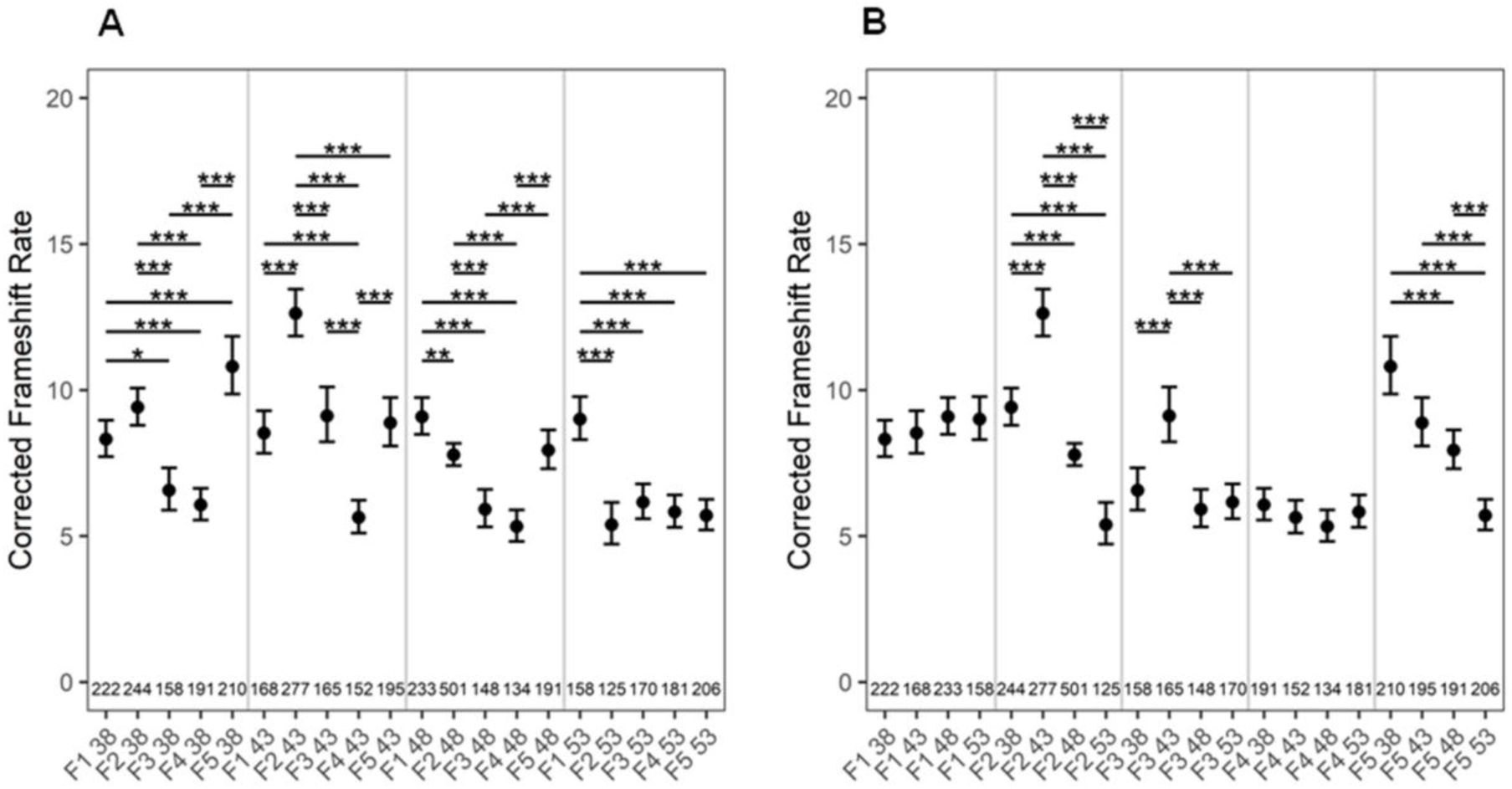
A-B: FS mutation rates as a function of A. fixed, B. increasing parental age across generations. The numbers at the bottom of the graph show the number of seedlings analyzed. *P* values were corrected for multiple testing. No asterisk denotes no significant difference. *, *P <* 0.05; **, *P* < 0.01; ***, *P* < 0.001.

### Wide fluctuation in transposition rates across same or different ages/ generations

To score transposition rates as a function of constant/advancing parental age across different generations, transgenic *Arabidopsis* plants containing the transposable element *tag1* between a 35S promoter of Cauliflower mosaic virus (CaMV) and the *uidA* gene were used (Liu and Crawford, 1998). *tag1* excision allows the expression of *uidA* gene, resulting in blue colored spots upon a histochemical assay. Although transposition rates as a function of constant parental age exhibited a pattern of a increase succeeded by a decrease, the differences were insignificant, with few exceptions (Fig. 8A). The transposition rates were random compared to other classes of mutations (Fig. 8A, B).

**Figure 8.**
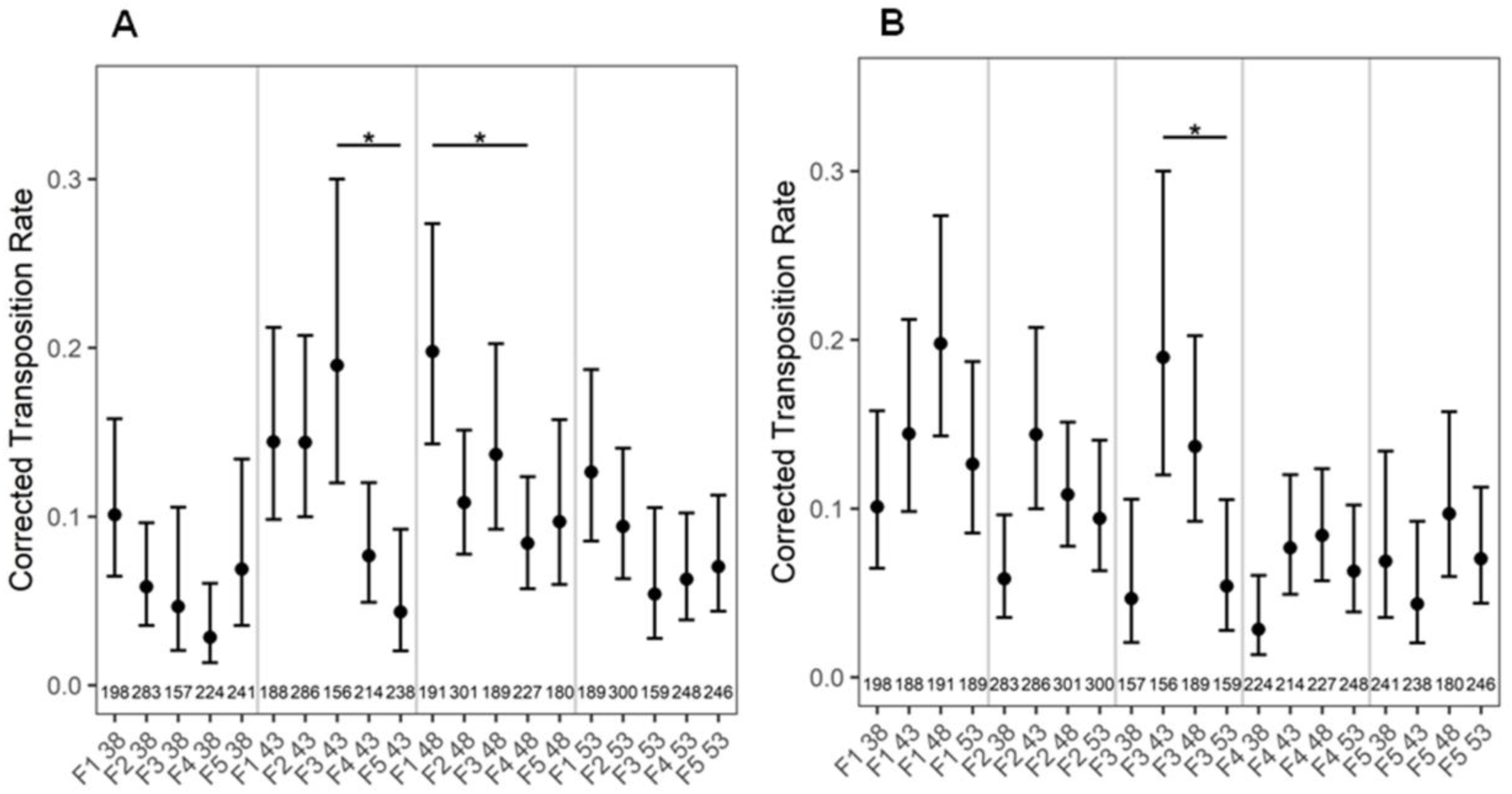
A-B: Transposition rates as a function of A. constant B. increasing parental age across different generations. The numbers at the bottom of the graph show the number of seedlings analyzed. *P* values were corrected for multiple testing. No asterisk represents no significant difference. *, *P <* 0.05.

### Irrespective of age, reversion rates fluctuate widely in seedlings from fourth and fifth generations

We further wanted to determine if the shift in mutation rates was significant within a certain generation, independent of age or conversely, if a significant change in reversions occur at a certain age regardless of the generation. For this, we combined mutation rates across all ages in a particular generation (for a particular class of mutation) and independently combined the reversion rates across all five generations within a particular age for a particular class of mutation. Irrespective of parental age, BSR, ICR, and FS rates followed similar trends with a small increase in the 2^nd^, a drop in the 3^rd^, followed by a significant reduction in the 4^th^ and a subsequent increase in the 5^th^ generation (Fig. 9 A, B, and C). Transposition rates also decreased significantly in the 4^th^ generation, however a similar surge in 5^th^ generation was not observed (Fig. 9D), reiterating our previous findings where all other classes of somatic mutations rates except transpositions exhibited an increase in the 5^th^ generation.

**Figure 9.**
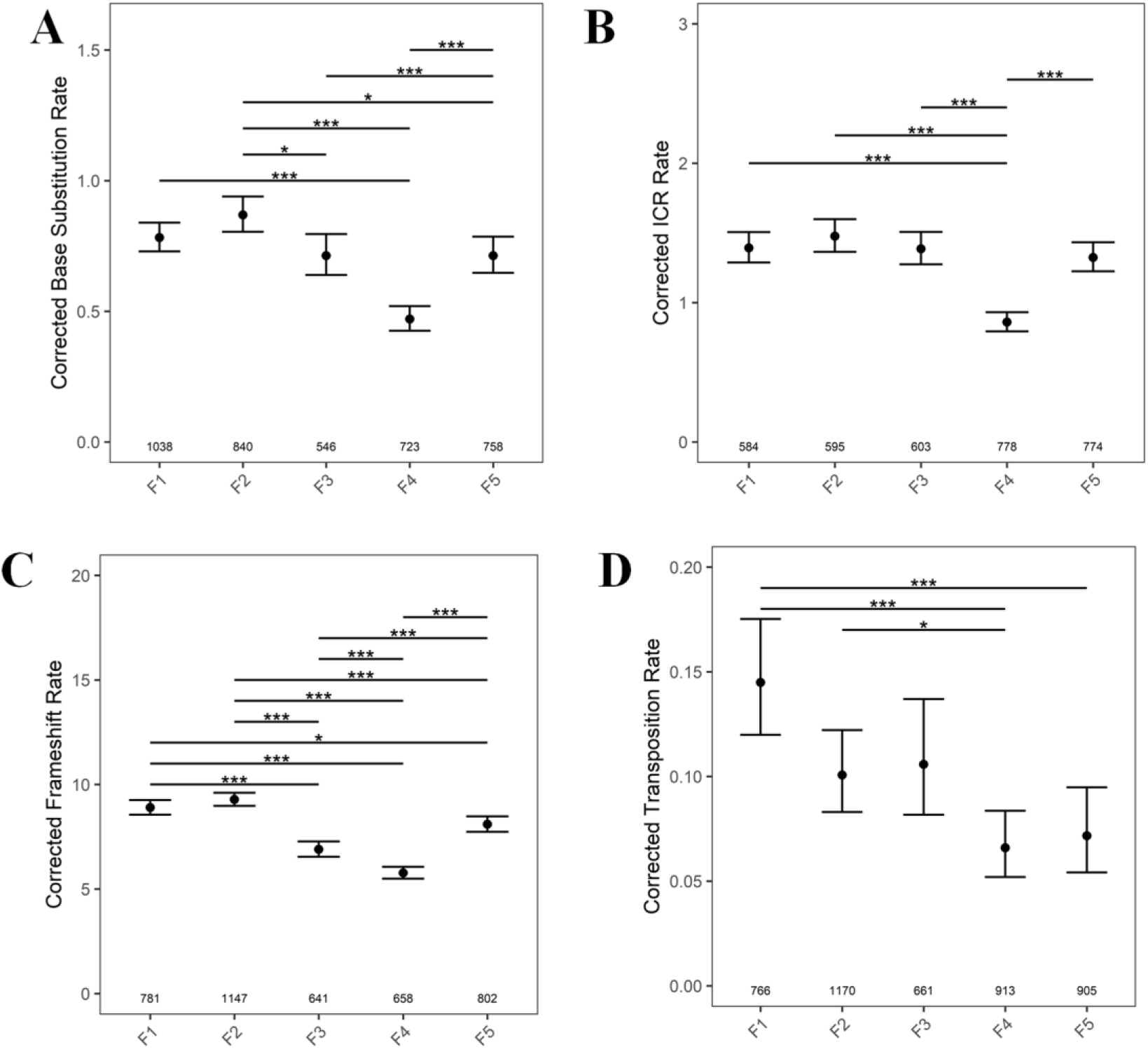
Sum total of mutation rates across generations (A) BSR, (B) ICR, (C) FS rates (D) Transposition rates in F1-F5 generation. The numbers along the X axis, show the numers of seedlings analyzed. *P* values were corrected for multiple testing. No asterisk represent no significant difference. *, *P* < 0.05; ***, *P* < 0.001.

Regardless of parental age, the average ploidy, the cell number, and cell size also show a significant change in the 4^th^ generation. Average ploidy is highest in 2^nd^, significantly decreased in 4^th^ and surged in 5^th^ generation (Fig. 10A). The observed leaf surface area (Fig. 10C) and adaxial epidermal cell number (Fig. 10D) were highest in the 4^th^ and reduced drastically in the 5^th^ generation. In contrast, there was no change in cell size in the first three generations, but dropped significantly in 4^th^ and increased thereafter in the 5^th^ generation.

**Figure 10.**
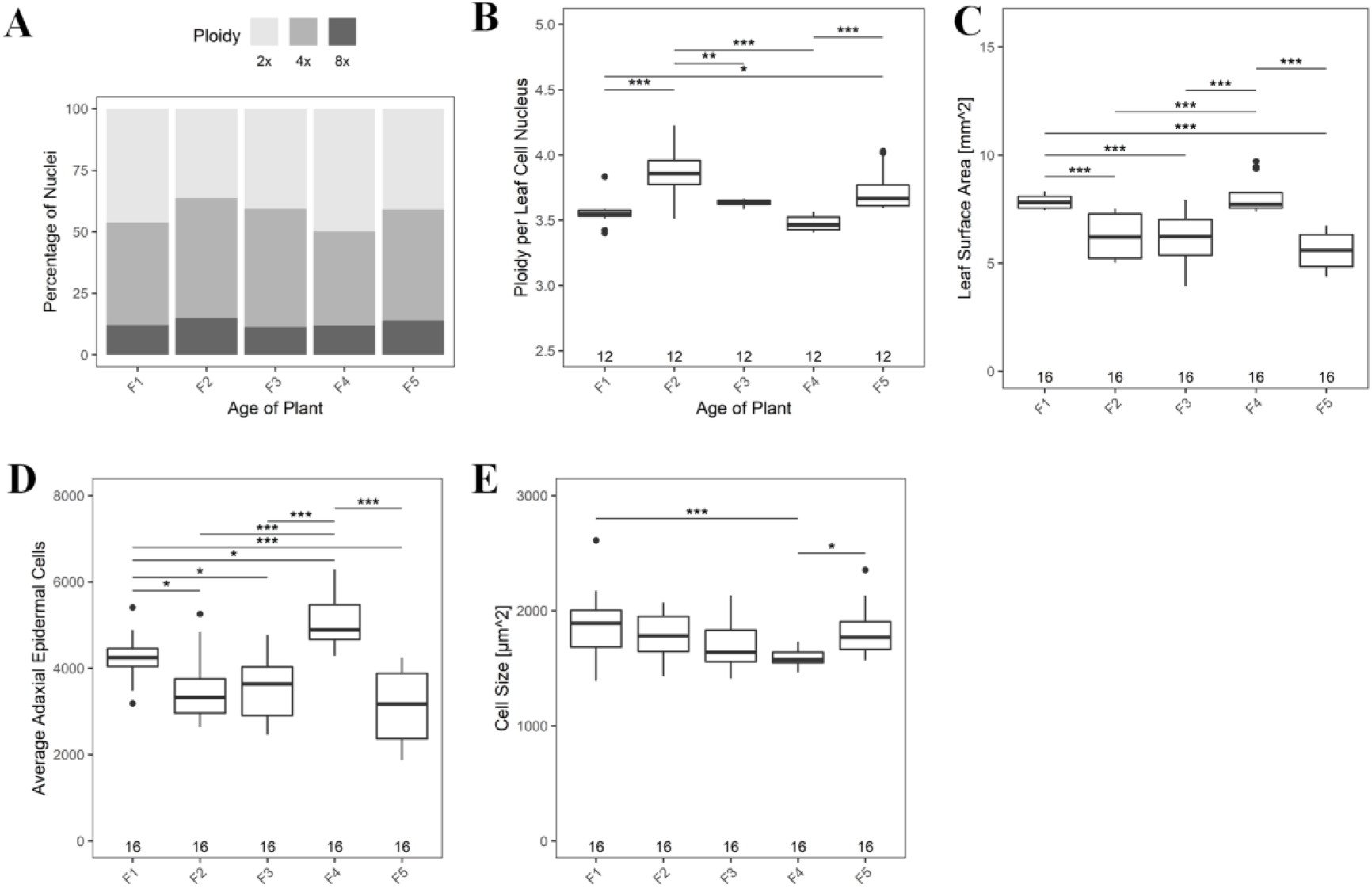
The graph representing the generation wise comparison of (A) total percentage of 2X, 4X and 8X nuclei, (B) Average ploidy per leaf per cell nucleus. (C) Total leaf surface area. (D) Average adaxial epidermal cell numbers. (E) Cell size of leaf. No asterisk represent no significant difference. *, *P <* 0.05; **, *P* < 0.01; ***, *P* < 0.001.

We then examined if mutation rates changed as a function of parental age independent of the generation by pooling mutation rates across all generations within a particular age. Although, ICR, FS, and transposition rates followed similar trends, with rates increasing in seedlings of 43 DAS, the subsequent decline in FS rates was apparent only from seedlings of 48 DAS. However, the significant drop in ICR, FS, and transposition rates was only in seedlings of 53 DAS plants and not from other ages (Fig. 11 B, C and D).

**Figure 11.**
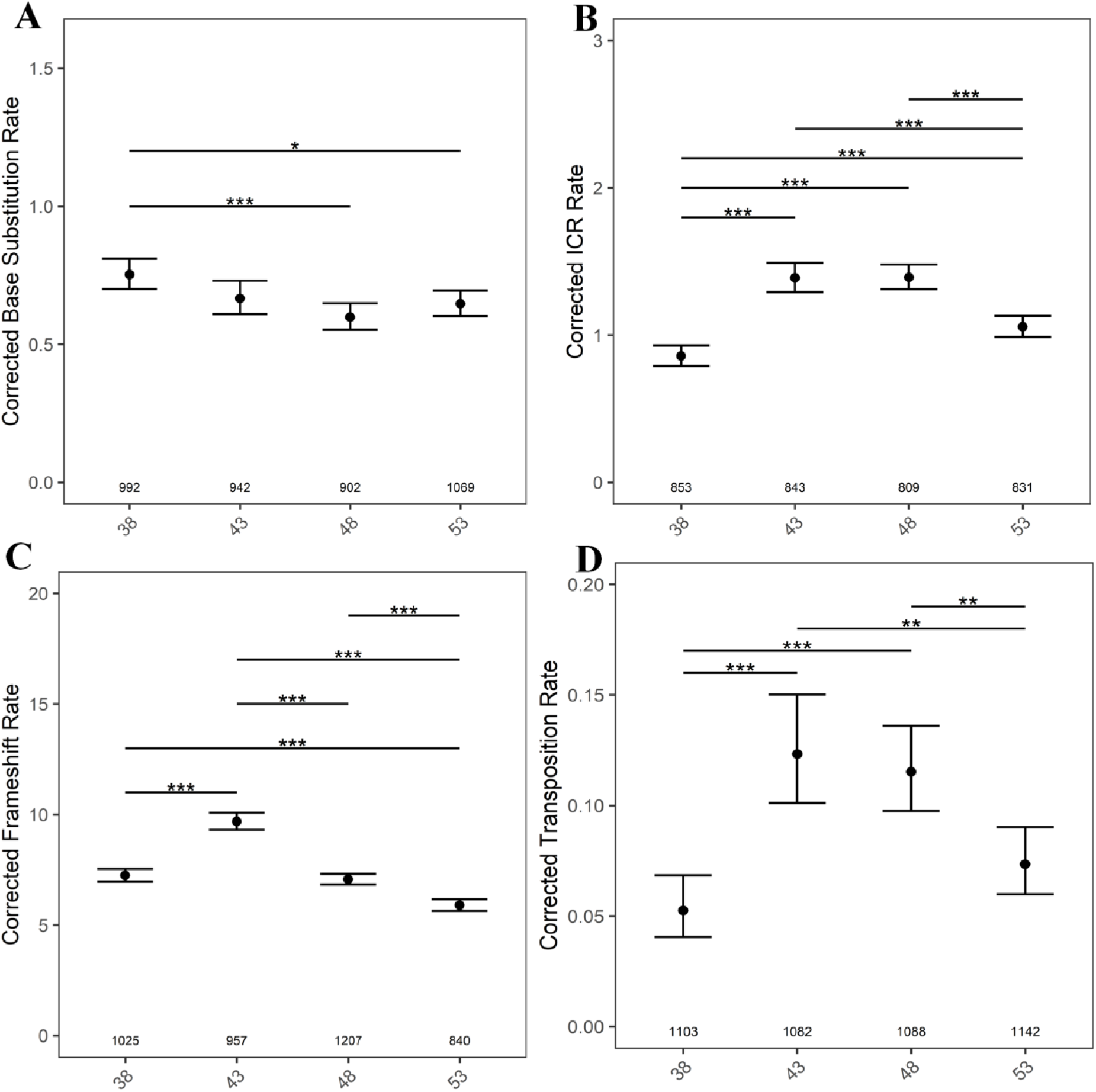
The graph representing the age-wise comparison after combining all five generations. (A) BSR, (B) ICR, (C) FS rates (D) Transposition rates in seedlings from 38, 43, 48 and 53 DAS plants. The numbers along the X axis, show the number of seedlings analyzed. *P* values were corrected for multiple testing. No asterisk represent no significant difference. *, *P* < 0.05; ***, *P* < 0.001.

Regardless of the generation, no significant variation was observed in the average ploidy but cell size decreased gradually with parental age (Fig. 12E).

**Figure 12.**
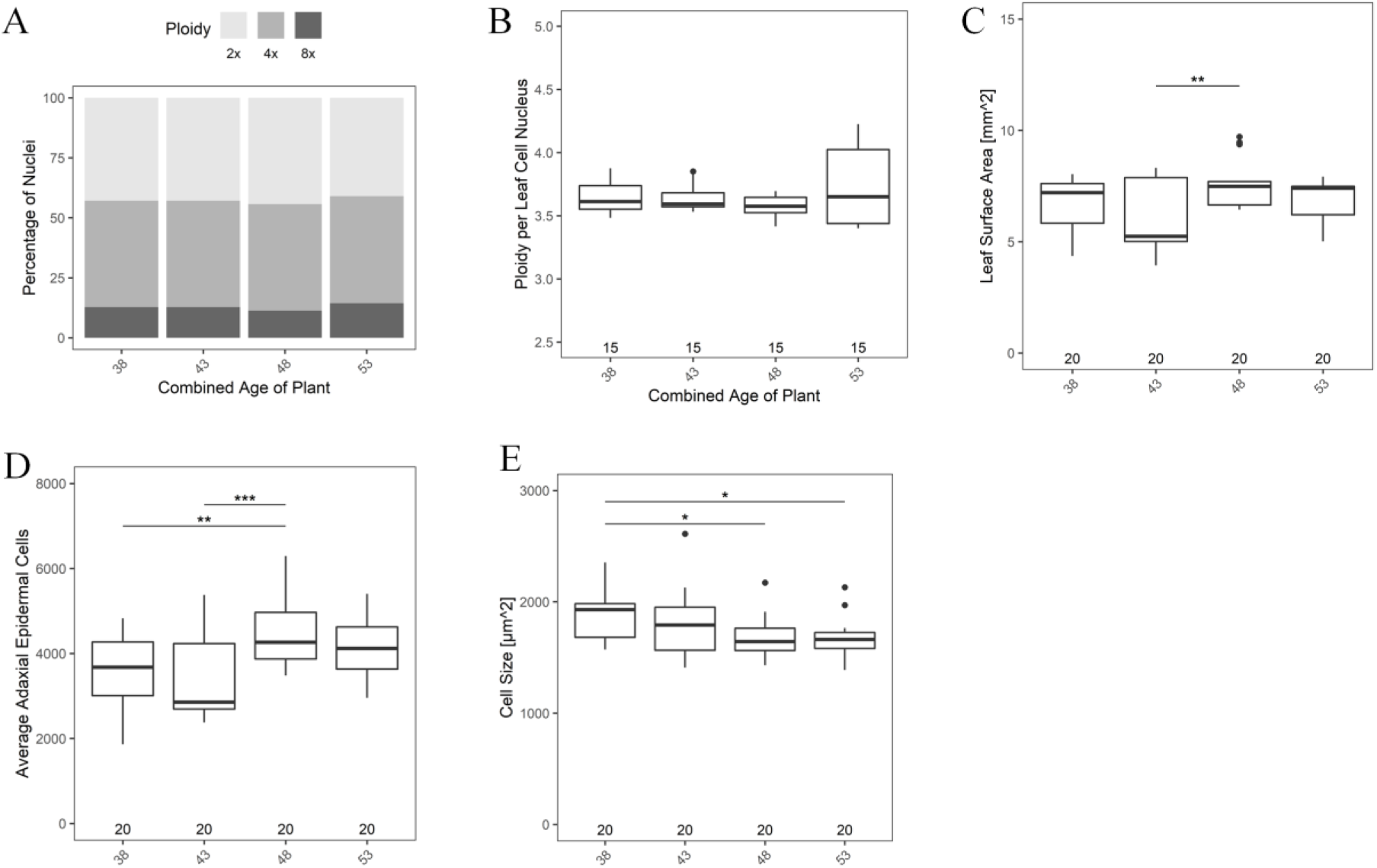
Graphs representing the age-wise comparison of ploidy, cell number, and cell size from all five generations. (A) Percentage of 2X, 4X, and 8X nuclei. (B) Average ploidy per leaf per cell nucleus. (C) Total leaf surface area. (D) Average adaxial epidermal cell number (E) Cell size of leaf.. No asterisk represents, no significant difference. *, *P <* 0.05; **, *P* < 0.01; ***, *P* < 0.001.

### Double strand DNA damage increases with increasing parental age in all generations

As age is known to increase the mutation rates, we employed a COMET assay to determine if double-strand DNA (ds-DNA) breaks increase in seedlings derived from parents of different ages. In this assay, damaged DNA forms a comet-like structure during migration in an electrophoresis gel (Moorad and Promislow, 2008). ds-DNA breaks were found to increase gradually in seedlings from advanced parental ages, reaching a maximum from seedlings of 53 DAS plants. The pattern of DNA double-strand breaks were similar (Fig. 13A) in the 1^st^, 2^nd^, and 5^th^ generations and again between 3^rd^ and 4^th^ generations. When parental age across generations was retained, no significant change in DNA damage was observed (Fig. 13B).

**Figure 13.**
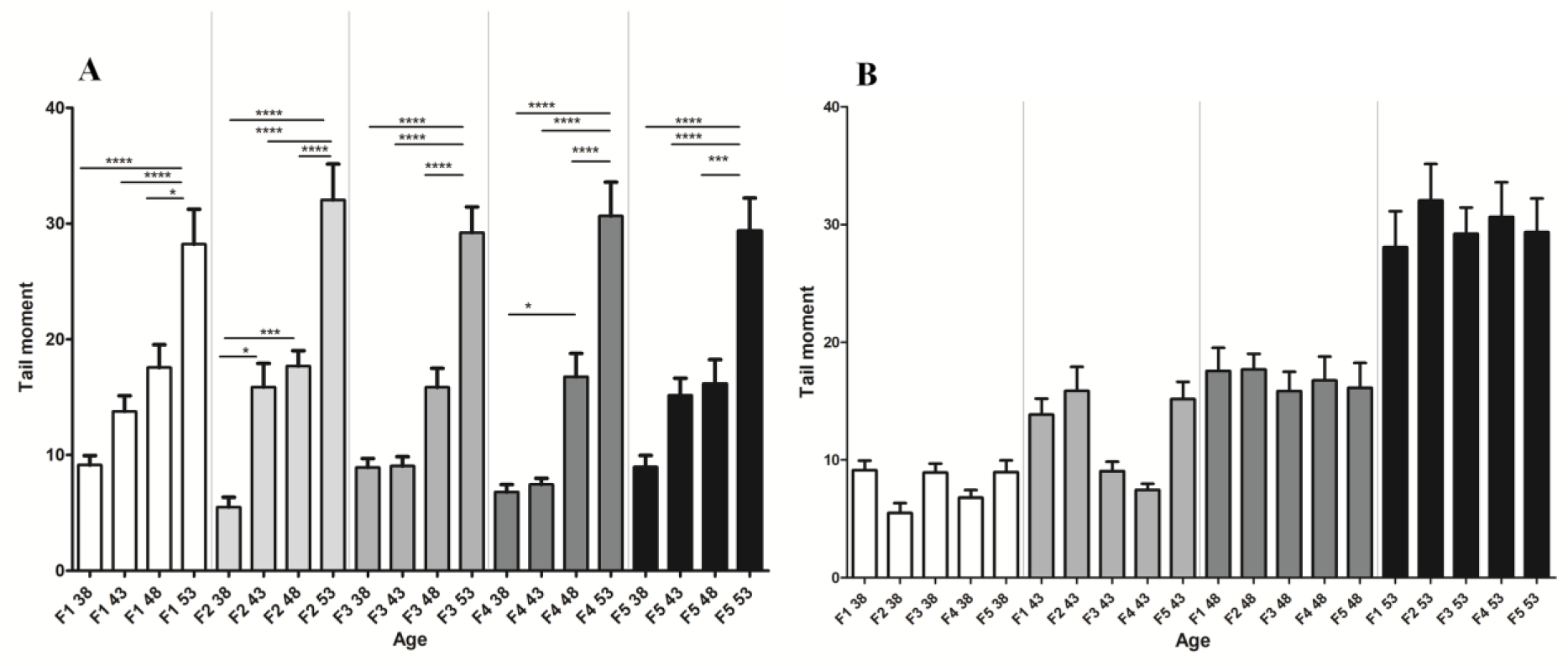
The DNA double-strand break as measured by a COMET assay. (A) Represents the DNA double-strand break with respect to increasing parental age in different generations. (B) Represents the DNA double-strand break with respect to fixed parental age in different generations. No asterisk denotes no significant differences *, *P **<*** **0.05; ***, *P* < 0.001; ****, *P* < 0.0001.**

### Wide variation in the expression levels of genes involved in DNA repair in first four generations but relatively stable expression in the 5^th^ generation

We examined if there exists a correlation between somatic mutation rates and expression of candidate genes involved in DNA repair such as *ATM, BRCA1, RAD 51* (Friesner *et al.,* 2005; Bleuyard *et al.,* 2005). Methylation levels are also known to impact somatic mutations (Graaf *et al.,* 2015), and hence, the expression of candidates genes like *DDM1, MET1* (Tan *et al.,* 2018; Zubko *et al.,* 2012) were also analyzed together with *5.8S rRNA* (McStay and Grummt, 2008; Simon *et al.,* 2018) by quantitative real-time PCR.

Although few interesting patterns in gene expression across different ages/generations were observed, this did not correlate with mutation rates. For instance, in seedlings obtained from 48 DAS plants, *RAD51* expression increased in the 4^th^ and dropped in the 5^th^ generation, while in contrast, seedlings from 53 DAS plants exhibited low expression in the 2^nd^ and 4^th^ generation, but increased significantly in the 5^th^ generation (Fig. 14A). With increasing parental age, *RAD51* expression was significantly upregulated in seedlings from 53 DAS plants from the 3^rd^ and 5^th^ generation compared to F1 38 DAS plants (Fig. 14B) but we found no correlation between the expression pattern and the observed mutation rates. Although *ATM* expression was consistently down-regulated, the drop in expression was significant in the 2^nd^ and the 5^th^ generation in all age groups (Fig. 14C). However, with increasing parental age, *ATM* was down-regulated significantly in 2^nd^, 3^rd,^ and 5^th^ generations, compared to F1 38 (Fig. 14D). *BRCA1* expression was significantly down-regulated across all ages (Fig. 14E) and with advanced parental ages, this pattern was observed only in the 4^th^ (2.7 fold) and 5^th^ (2.51 fold) generations compared to 38 DAS of the 1^st^ generation (Fig. 14F). While there was considerable heterogeneity in *MET1* expression levels across all ages/generation, the fold change (7.74 to 23.4) of *MET1* expression was striking across all ages in the 5^th^ generation alone (Fig. 14G). Plants of advancing ages (53 DAS) bred at the same age for five generations show an interesting *DDM1* expression pattern of an initial high, drop in 2nd, gradual increase thereafter, and another drop in the 5^th^ generation. Except for a few, there was no change in *DDM1* expression as a function of increasing parental age (Fig. 14J). *sRNA* shows contrasting expression patterns depending on the parental age. For instance, in 38 DAS, the expression was high initially then dropped for the latter four generations, while in older (53 DAS) plants, the expression was initially low in the 1^st^ generation, and underwent an increase in subsequent generations (Fig. 14K). The wide variation in gene expression levels of different candidates analysed in the first four generations and relatively stable expression in the 5^th^ generation may have a combinatorial effect on mutation rates, which requires further investigation.

**Figure 14.**
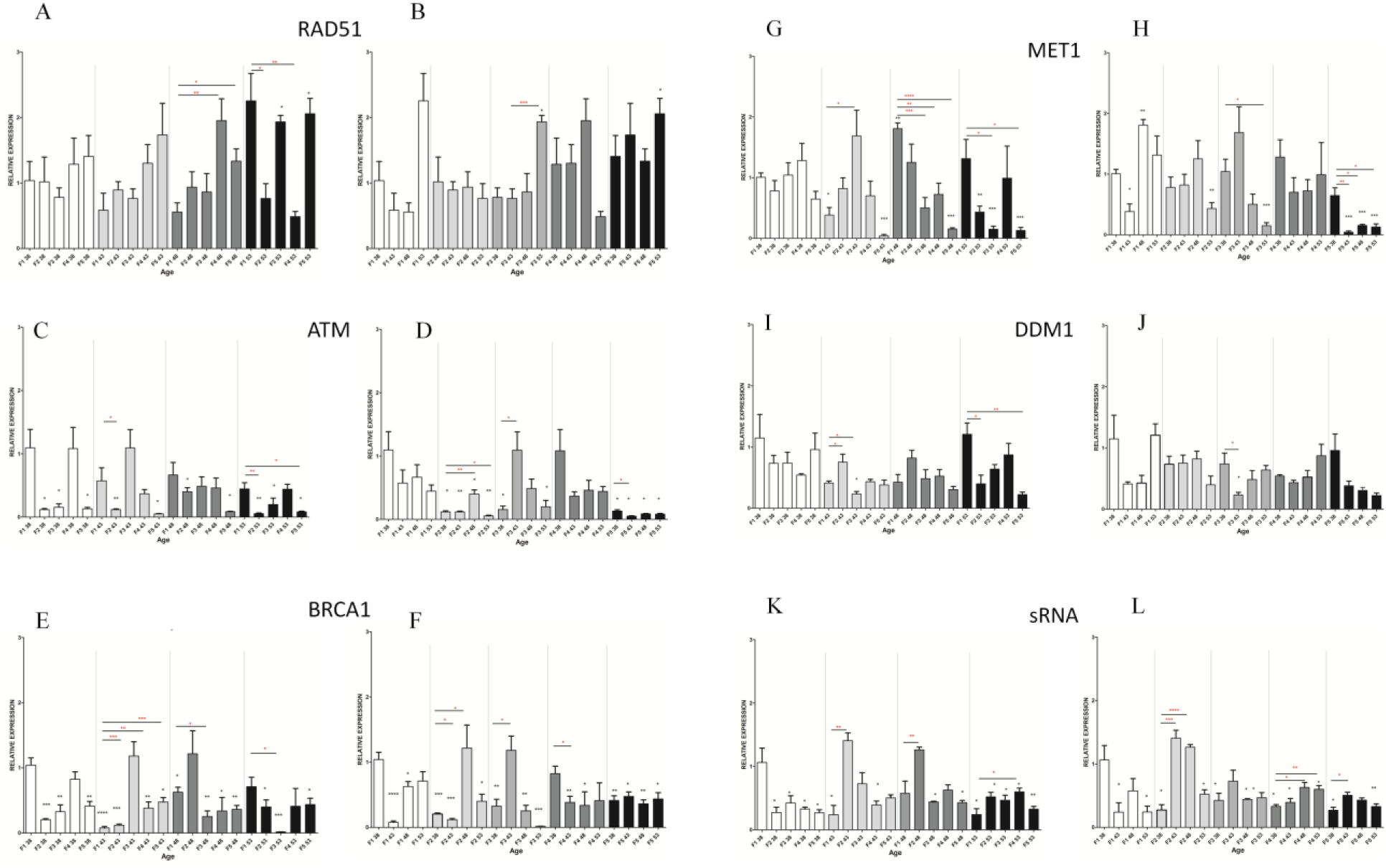
The relative expression of *RAD51* (A), *ATM* (C), *BRACA1* (E), *MET1* (G), *DDM1* (I), small RNA 5.8 (K). The black star represents a significant change in expression in comparison to F1 38 (*) and the significance between same age groups /generations are represented by a red star (*). The graph on the left panel represents gene expression analysis from same-aged parents across different generations. The graph showing the relative gene expression of *RAD51* (B), *ATM1* (D), *BRACA1* (F), *MET1* (H), *DDM1* (J), *small RNA 5.8* (L) representing comparative gene expression with increasing parental age in every generation (with *GAPDH* as a reference gene). Error bars represent SEM of biological replicates (n =3–6). Student’s *t*-Test *, *P <* 0.05; **, *P* < 0.01; ***, *P* < 0.001; ****, *P* < 0.0001.

## Discussion

### Possible reasons for the differences in the mutation rates between the work of Singh *et al.,* (2015) and this study

Using similar set of detector lines, (Singh *et al.,* 2015) reported mutation rates that were considerably different from this study, although there were some similarities in the patterns. Several reasons could account for the observed inconsistencies between the two data sets. Previously, (Singh *et al.,* 2015) used separate plants to ascertain the paretal age effect that involved emasculation followed by manual self-pollination. However, throughout this study flowers fertilized on different DAS were identified in a single plant without any emasculation. Emasculation involves, holding and removal of accessory flowers from the inflorescence. Given that physical damage or repeated mechanical stress is known to elevate the mutation rates (Pastor *et al.,* 2013; Heyer *et al.,* 2018; Liu *et al.,* 2021 Molinier *et al.,*, 2006), emasculation may also increase the somatic mutation rates. Further, manual self pollination was carried out after two days after emasculation and the stress reponse is known to significantly depend on plant age (Saini *et al.,*, 2017) which would bias the mutation rates. The detector lines are extremely sensitive to small environmental perturbations and even a brief variation in growth temperature resulted in increased mutation rates. For example, the FS rates goes up by ∼5 fold after increasing the temperature from 12^0^C to 22^0^C (Saini *et al.,*, 2017). Moreover, housing such detector lines in proximity to plants subjected to UV stress is known to trigger genome-wide changes and elevated mutation rates, likely due to the effect of volatiles from treated plants (Yao *et al.,*, 2011). Hence to avoid emasculation-induced stress responses that would potentially bias mutation rates even within control plants nearby, we identified and tagged flowers which just undwerwent pollination at various DAS, following which the seeds were harvested. Furthermore, unlike the previous study, we took into account the influence of seed age, and harvested the seeds and plated them immediately at maturity. Ideally, storing the seeds from plants of different ages/generations and staining all the seedlings at the same time will remove any bias from the resultant data, due to uniform variation in staining across all seedlings. However, this approach would also strongly bias the results since seed storage is a strong determinant of mutation rates, even upon storage at 4^0^C (Thakur *et al.,* data not published). In the earlier work of (Singh *et al.,* 2015), for each parental age, emasculation was carried out from a set of flowers from independent plants and in contrast, we identified all the self-pollinated flowers of a particular age following anthesis from the same plant and this could also bias mutation rates. However, extreme care was taken to avoid even small environmental perturbances. Every set including control plants were grown in different chambers in different compartments in a highly randomized fashion and the physical position was changed every four days. Thus, the collective effect of the individual variables may account for the observed differences in mutation rates.

Further, Although we found similar pattern of BSR, ICR, FS, and transposition rate as a function of parental age in manual (Fig. 4 A, B, C, D) and self pollinated studies (Fig. 4 E, F, G, H), there is variation in ICR rates between this work (Fig. 4) and that of (Singh *et al.,* 2015). Earlier, (Singh *et al.,* 2015) used the detector lines R2L1 and R3L30 to score ICR rates and found that parental age had little impact on ICR rates in R2L1, while in R3L30, ICR rates declined with age. In contrast, we observed an increase in ICR rates with parental age in R2L1 line. Several reasons could account for the observed differences, including the number of generations the plant has passed through selfing and the seed age.

Using such detector lines, independent studies from the same lab observed different reversion rates just in controls alone (Ilnytskyy *et al.,* 2004; Boyko *et al.,* 2005, 2006). For example, line 11 reporting homologous recombination show a reversion rate of 1.9, 2.71 and 2.5 whereas in line 651, the reversion frequency was found to be 0.21, 0.74, and 0.7 across three different studies. All these three studies involved using homozygous detector lines and seedlings of the same ages were used to score GUS reversions. Similarly, detector line IC1 and IC9 reporting ICR were tested, and independent observations from the same lab found different reversion rates for particular line (Molinier *et al.,* 2004, 2006). In IC1, the recombination frequency in controls were 0.32 and 0.09, whereas in line IC9, the values were 0.18 and 0.04 in controls. Thus for a particular line, the number of reversion events were not consistent across studies and possibly distorted due to a variety of factors including seed storage age, parental age and the number of generations the plants carrying GUS has gone through selfing at a particular DAS. In light of these factors, care was taken to avoid seed storage and seeds were plated soon after harvesting. However, the seeds were obtained from different labs at different times and the number of times the plants has gone through selfing subsequent to transformation is unknown. In plants, the persitance of stress memory effect is known to fade away after four generations (Molinier *et al.,*, 2006). The detector lines were maintained by selfing and thus the effect we observed is likely not due to stress or small environmental perturbances, but from parnetal age differences alone. Furthermore, the replicates for the GUS assay were grown at different time but within the same chambers and thus the role of environemtal perturbation in modulating the mutation rates was minimized. By comparing the whole genome sequence of *A. thaliana* derived from a single seed descent, somatic mutation rates were estimated in two widely separated generations. In *A. thaliana*, somatic mutations were found to be biased toward G:C to A:T and are concentrated in transposable elements, pericentrometic/centromeric zones (Weng *et al.,* 2019). However, it was not known if the fertilization age was retained across successive generations in that study. Although such an approach is insightful, there are limitations. Since reversion events are rare and occur randomly in seedlings, sequencing approaches may not be optimal for detection. Our results strongly suggest that somatic mutation rates follow a specific pattern that significantly depends on the number of generations a plant has gone through selfing at a particular age. Although the number of generations we analysed is small, the data strongly indicates that somatic mutation rates follow a pattern, robustly influenced by function of parental age and generation time. This data suggests that parental age alone contributes to fluctuations in mutation rates, with different ages exhibiting different patterns of fluctuations, however the molecular mechanisms underlying this phenomenon is unknown. Such stochastic responses are a part of successful evolutionary adaptations in unprediactable and everchanging environments (Shahrezaei and Swain, 2008).

The expression pattern of genes involved in DNA repair such as *RAD51*, *ATM,* and *BRCA1* and *sRNA* was random in the first four generations but became relatively stable in the 5^th^ generation (Fig. 14). The number of generations may be counted by fluctuations followed by stable gene expression impacting the mutation rates as in every 5^th^ generation there is a surge in C to T transition rates. In *C. elegans*, for example, changes in histone methylation expression are stochastic for the first four generations, and thereafter becames stable in the 5^th^ (Greer *et al.,* 2011). If such changes are conserved in *Arabidopsis* also, it would likely have an impact on gene expression and mutation rates. An increase in C→T base substitution events is related to cytosine methylation, and methylcytosine is susceptible to deamination (Mugal and Ellegren, 2011). High degree of DNA methylation, possibly leads to low gene expression (Moore *et al.,* 2013) and such genes transcribing at reduced levels are likely to have low mutation rates as well (Park *et al.,* 2012). DNA methylation levels are known to be higher in older compared to younger plant tissues (Bitonti, 2002; Fraga *et al.,* 2002; Sun *et al.,* 2015) but some studies have also reported contradictory findings (Monteuuis *et al.,* 2009; Meng *et al.,* 2012; Michalak *et al.,* 2015). Cytosine DNA methylation is important for maintaining genome stability and regulating gene expression (Vanyushin and Ashapkin, 2011) but is also known to increase neighborhood mutations in somatic cells of *Arabidopsis* (Kusmartsev *et al.,* 2020). This suggests a context-dependent methylation event during development (Ambrosi *et al.,* 2017; Baker *et al.,* 2018). Our results show higher C→T transition rates in seedlings obtained from younger and older, but not from middle-aged parents. *DDM1* and *MET1* are required for CG methylation. In support of a role for *DDMI* in higher C→T transitation from younger and older parents, a correlation was observed between *DDM1* expression (Fig. 14J) and mutation rates in seedlings from middle-aged parents (43 and 48 DAS) of 1^st^ and 3^rd^ generations and similar mutation rates in 4^th^ and 5^th^ generation (Fig. 5B). Our results show increased ICR events in seedlings of 38, 43, and 48 DAS plants, but not from seedlings of 53 DAS plants (Fig. 6B). Although DSBs increased with parental age, we still observed low ICR rates in seedlings obtained from 53 DAS plants, which may be due to increased non-homologous end-joining (NHEJ) with plant age (Boyko *et al.,* 2006). The expression of Ku70 protein in particular, known to be involved in NHEJ was identified to increase with plant age (Golubov *et al.,* 2010). This state resulting in a decline within ICR rates in older parents may be transmitted to the next generation, which would also explain the observed increase in DSBs in the offspring of older parents.

Plants with long life may have low cell division rates and, as a consequence, are also possess low mutation rates (Watson *et al.,* 2016). It is also possible that the cell division rates in the reproductive meristem may not be uniform across different ages. Furthermore, DNA replication is known to be independent of life span, suggesting that older plants may not be passing on more mutations to their offsprings relative to younger plants (Watson *et al.,* 2016). It is known that age-related accumulation of somatic mutations results in inactivatation of random genes critical for somatic cell function (Failla, 1958; Szilard, 1958; Moorad and Promislow, 2008). Therefore, epigenetic inheritance, possibly mediated by DNA methylation or small RNAs, may be a key mechanism underlying the observed transgenerational effects (Herman and Sultan, 2011; Holeski *et al.,* 2012; Zhang *et al.,* 2013; Amtmann *et al.,* 2013). Further, it is already reported the transgenerational stress memory effect tends to lost after 4 generation (Molinier *et al.,*, 2006) and considering age as a kind of stress which is responsible for the decrease of mutation rate till 4 generation but in 5^th^ generation its effect is lost and thus mutation rate increases in 5 ^th^ generation.

## Material and methods

### Plant growth conditions

Prior to planting, *Arabidopsis thaliana* (Columbia) seeds were surface sterilized with 70% (v/v) ethanol followed by 0.5% (v/v) bleach treatment for 3 min. The seeds were washed thrice with sterile water and plated on autoclaved MS medium (with 3% [w/v] Suc), pH 5.7, and containing 0.05% (v/v) Plant Preservative Mixture (Biogenuix Medsystem Pvt. Ltd.) and kept at 4°C for 48 h in the dark for synchronized seed germination. The plates were transferred to plant growth chambers (Percival, USA) that had a uniform light intensity of 8,000 lux (under a 16-h-light/8-h-dark cycle). The temperature of the growth chamber (Percival CU-36L6) was maintained at 22°C throughout, and the humidity was set to 80%. Three-week-old seedlings were transferred from MS plates to soil and grown again in a growth chamber (Percival AR-36L3). The soil consisted of equal proportions of garden soil, peat, perlite, and vermiculite. Plants were grown in a randomized manner in the growth chambers and at frequent intervals, their positions were changed.

### Somatic mutation detector lines

All the detector lines used in this work are in Col-0 background. The base substitution detector line 1,390_T→C_ was a gift from Anna Depicker, University Ghent University, Ghent, Belgium (Auwera *et al.,* 2008). Transgenic ICR lines (R2L1) carrying inverted catalase introns (418 bp) in the *uidA* gene and the frameshift detector line (G10) were a gift from Francois Belzile, Université Laval, Quebec, Canada (Li *et al.,* 2004; Azaiez *et al.,* 2006).

The detector line harboring the *Tag1* element was a gift from Nigel M. Crawford, University of California, San Diego, USA (Liu and Crawford, 1998).

### Identification of self pollinated flowers from plants of different DAS

Self-pollinated flowers were identified on 38, 43, 48, and 53 DAS from the same plant and marked with different colored threads after anthesis. Matured seeds were harvested and germinated on MS plates.

### Histochemical staining for GUS activity

GUS staining was performed in 3-week-old seedlings as described by Jefferson *et al.,* (1987) in three biological replicates. The total number of seedlings analyzed is indicated in the Figures. The blue colored GUS reversions were counted using a stereozoom microscope (Leica KL300, Germany).

### Estimating correction factors to calculate mutation rates

The mutation rates were not corrected for replication cycle but for the genome number. Mutation rates were calculated by dividing the average number of GUS spots per plant by the copy number of the transgene (Kovalchuk *et al.,* 2000). Since the total number of cells and the average ploidy per nucleus differ between seedlings from different generations, the total genome number will not be the same also. Hence, mutation rates were corrected by accounting for changes in the number of adaxial epidermal cells of the fourth true leaf and the average ploidy in 3-week-old seedlings (derived from F2, F3, F4 and F5 generations of 43, 48, and 53 DAS). This was compared with the adaxial epidermal cells of the fourth true leaf and ploidy levels in seedlings derived from F1 38 DAS plants. The correction factor was calculated using the formula (Singh *et al.,* 2015).

TITER = (P_H_ ×C_H_)/ P_Y_×C_Y_

To compare differences in mutation rates as a function of plant age, seedlings derived from young parents (F1 38 DAS) were considered (Singh *et al.,* 2015). *P*_H_ is the average ploidy per nucleus in 3-week-old seedlings from 43, 48, and 53 DAS plants of F2, F3, F4, and F5 generation. *C*_H_ is the average number of adaxial epidermal cells in the fourth true leaf of seedlings derived from 43, 48, and 53 DAS plants of F2, F3, F4, and F5 generation. *P*_Y_ is the average ploidy per nucleus in 3-week-old seedlings from F1 38 DAS. *C*_Y_ is the average number of adaxial epidermal cells in the fourth true leaf of seedlings derived from F1 38 DAS plants.

Corrected mutation rate = GUS/titer

where GUS is the average number of blue spots per plant.

### Statistical analysis

A total of 3905, 3336, 4027, and 4415 seedlings were examined to detect BSR, ICR, FS, and transpositions, respectively. The revertant GUS spots are random and do not follow normal distribution and hence ANOVA test was avoided. The number of GUS spots are count data and to account for overdispersion in the data we chose a Quasi-Poisson generalized linear model (GLM) with the log link function (Nelder and Wedderburn, 1972). The log of the correction factor for cell number and ploidy per leaf cell nucleus was integrated into the models as a fixed intercept. In all GLMs, the data from the groups were used for multiple comparisons. Correction for multiple testing was done to keep the family-wise error rate at 5% (Gabriel, 1969). *P* values were adjusted with a single-step method that considers the joint multivariate *t* distribution of the individual test statistic (Bretz *et al.,* 2010). The results are presented with a two-sided *P* value adjusted for multiple comparisons. All the statistical analysis were done using R (R Developmental Core Team, 2010) For multiple comparisions, *P* values were modified by multcomp package R (Bretz *et al.,* 2010). Graphs were produced with ggplot2 (Wickham, 2009).

### Cell size and cell number analysis by scanning electron microscopy

For SEM, a wax impression of plant tissue was prepared according to the protocol of Beermann and Hülskamp (2010). The fourth true leaf of a 3-week old *Arabidopsis* seedling derived from parents of different ages was dissected, and as suggested by the manufacturer, the two components of waxy dental material were deposited on the leaf to generate an impression (Coltene PRESIDENT light body, Coltene AG, Altstatten, Switzerland). After 5 min, when the wax had hardened, the leaves were gently removed. This leaf mould was used for sample preparation with Spurr resin, and the resin with the leaf impression was taken out carefully from mould, and the sample was coated with gold using Polaron Range sputter coater (Quorum Technologies). The coated resins were mounted onto a SEM stub with a double-sided carbon tape to capture the images (FEI Quanta 400-F SEM (FEI Company, USA) under 20kV voltage and 70 Pa pressure). The adaxial leaf surface area was calculated by using the SEM images captured at 50X magnification (Singh *et al.,* 2015). Independently, SEM images were taken at different positions of the leaf at 500X magnification. The average cell size was calculated by dividing the number of cells observed in an area of 500X magnification at different positions of the leaf. To calculate the total number of adaxial epidermal cells, the total area of the leaf was divided by a fixed area of 500X magnification and multiplied by the number of cells present in an area of 500X magnification. ImageJ (NIH, USA) software was used for counting the total number of cells and the total area of the leaf. Four biological replicates were considered to determine the average number of adaxial epidermal cells of the 4^th^ true leaf.

### Ploidy analysis by flow cytometry

Ploidy analysis was performed as per the protocol of Doležel *et al.,* (2007). The fraction of nuclei with 2X, 4X, and 8X ploidy were estimated together with the average number of genomes per nucleus from seedlings of different parental ages. Approximately 60 mg of leaf tissue of 3-week-old seedlings was finely chopped with a razor blade in a glass Petri dish containing 1 mL of ice-cold nuclei extraction buffer (Sysmex CyStain^TM^ PI Absolute P KIT). The chopped tissues were filtered with a 20 μm nylon filter (CellTrics® filters, Sysmex). Three biological replicates were carried out for flow cytometry. To stain the nuclei, 2 ml of staining buffer (provided with the kit) containing propidium iodide and RNase in 1:2 ratio was mixed to the filtrate and kept for an hour at room temperature. A sample derived from Tomato (*Solanum lycopersicum* ‘Stupicke’) was used as control. After the incubation period, the samples were analyzed using a Partec CyFlow® Ploidy analyser (Sysmex). Data analysis was carried out with CyView Software (Sysmex).

### RNA isolation and quantitative real-time PCR

Using the Trizol reagent (Invitrogen, USA), total RNA was isolated from 21-day old seedlings of wild type Columbia-0 (Col-0) obtained from plants of different ages in five consecutive generations. cDNA was synthesized using 2 μg RNA, random hexamers, and High Capacity cDNA Reverse Transcription Kit (Applied Biosystems, USA). Quantitative real-time PCR was done in triplicate using the DyNAmoTM Flash SYBR Green qPCR kit (QuantStudio (TM) 7 Flex System). The primers for amplifying *ATM, BRCA1, RAD51, DDM1, MET1,* and *sRNA* (5.8S *rRNA* and 25S *rRNA* with 18S rRNA fragment) are listed in Table S1. The expression levels of glyceraldehyde-3-phosphate dehydrogenase (*GAPDH*) mRNA was used as an internal standard. The relative gene expression in each age and generation was determined by calculating 2^(−ΔΔCt)^ as described previously (Veremeyko *et al.,* 2012).

### Neutral COMET assay

To measure ds DNA breaks, a COMET assay was performed using the Oxiselect Comet Assay Kit from Cell Biolabs, Inc. USA. 3 week old Col-0 seedlings of 38, 43, 48, and 53 DAS from all five generations were chopped using a razor blade in 1 mL of Otto I solution. Subsequently, the sample was filtered with a 20µm nylon filter (CellTrics® filters, Sysmex), and the filtrate was centrifuged at 300 rpm for 5 min. The pellet was dissolved in phosphate-buffered saline (PBS) containing 20 mM EDTA. The dissolved sample was mixed with warm low-melting agarose in a ratio of 2:5 and poured onto a slide coated with normal agarose. The sample was covered with a glass coverslip, stored at 4°C for 15 min and after removing the coverslip, the slides were transferred to a chamber filled with chilled lysis buffer for 30 to 60 min at 4°C. The slides were then transferred to another chamber filled with cold alkaline solution (NaOH and EDTA, pH 10) for 30 min at 4°C in the dark. The slides with the sample were treated with chilled Tris-borate/EDTA buffer for 5 min and transferred to a horizontal electrophoresis chamber containing the same buffer. Electrophoresis was carried out at 1 V cm^-1^ for 15min. Thereafter, the slides were gently rinsed thrice with deionized water, followed by rinsing with 70% (v/v) ethanol for 5 min. After air drying, the sample was stained with vista green and kept for 15 min in dark at room temperature. Then, the COMETS were observed using an upright fluorescence microscope (Nikon Eclipse 80i) fitted with a fluorescein isothiocyanate (FITC) filter. We analysed around 50-55 Comet using a free online tools available from CaspLab. One way ANOVA with a 95% confidence interval of difference was used for the statistical analysis carried out by prism software (USA).

## Author contibution

Conceived and designed the experiments: SB and RB. Performed the experiments and compiled the data: SB and YT . Analysed the data: SB, AKS, and RB. Wrote the paper: SB, AKS, and RB. All authors read and approved the final manuscript.

## Acknowledgments

We gratefully acknowledge Francois Belzile (Université Laval), Anna Depicker (University of Ghent), Nigel M. Crawford (University of California, San Diego), and Barbara Hohn (Friedrich Miescher Institute) for providing reporter line seeds; The Sophisticated Analytical Instruments Facility (SAIF), Indian Institute of Technology-Madras for assistance with SEM analysis; and Rama Shanker Verma (Indian Institute of Technology-Madras, Chennai) for help with flow cytometry.

---

**Table S1.** The mutation rates were normalized by factoring the differences in cell number of the 4th true leaf and ploidy per nucleus from plants of different ages/generations. The correction factor was derived from a five-generation study of adaxial epidermal cell count and ploidy per nucleus of leaf cells. The Interquartile Range Coefficient (CIQR) was calculated as a non-parametric variance measurement in the style of the variance coefficient. The relative number of cells and the relative ploidy are the normalization values for generations F2, F3, F4, and F5 compared to F1, and for older ages (43, 48, and 53 DAS), the comparison was with 38 DAS plants. The correction factor was determined by combining the two values for standardization and was introduced before correcting the number of GUS spots. ‘n’ is the number of plants analyzed and median is the median of measurements.

| Epidermal Cells |  |  |  |  | Ploidy per Nucleus |  |  |  |  |
| --- | --- | --- | --- | --- | --- | --- | --- | --- | --- |
| Age | n | Median | CIQR | Relative Cell No. | n | Median | CIQR | Relative Ploidy | Correction Factor |
| F1 38 | 4 | 4134.841645 | 0.074743099 | 1 | 3 | 3.557 | 0.0402 | 1 | 1 |
| F1 43 | 4 | 4092.023519 | 0.111139791 | 0.989644555 | 3 | 3.577 | 0.0056 | 1.005622716 | 0.995209 |
| F1 48 | 4 | 4175.036919 | 0.119608416 | 1.009721116 | 3 | 3.544 | 0.0051 | 0.996345235 | 1.006031 |
| F1 53 | 4 | 4673.600815 | 0.120248471 | 1.130297413 | 3 | 3.425 | 0.0158 | 0.962890076 | 1.088352 |
| F2 38 | 4 | 3686.811979 | 0.025377695 | 0.891645266 | 3 | 3.866 | 0.0013 | 1.086870959 | 0.969103 |
| F2 43 | 4 | 2791.664624 | 0.097476684 | 0.675156358 | 3 | 3.841 | 0.0021 | 1.079842564 | 0.729063 |
| F2 48 | 4 | 4565.716472 | 0.165895059 | 1.104205884 | 3 | 3.558 | 0.0121 | 1.000281136 | 1.104516 |
| F2 53 | 4 | 3091.480092 | 0.081892202 | 0.747665898 | 3 | 4.209 | 0.0026 | 1.183300534 | 0.884713 |
| F3 38 | 4 | 3310.743566 | 0.177374671 | 0.800694162 | 3 | 3.639 | 0.0036 | 1.023053135 | 0.819153 |
| F3 43 | 4 | 2628.679586 | 0.078037259 | 0.635738877 | 3 | 3.594 | 0.0053 | 1.010402024 | 0.642352 |
| F3 48 | 4 | 4049.544556 | 0.109844988 | 0.979371135 | 3 | 3.654 | 0.0027 | 1.027270171 | 1.006079 |
| F3 53 | 4 | 4223.011129 | 0.160442252 | 1.021323545 | 3 | 3.651 | 0.0044 | 1.026426764 | 1.048314 |
| F4 38 | 4 | 4650.910075 | 0.02811808 | 1.12480972 | 3 | 3.486 | 0.0055 | 0.980039359 | 1.102358 |
| F4 43 | 4 | 5036.341781 | 0.046995477 | 1.218025311 | 3 | 3.556 | 0.0048 | 0.999718864 | 1.217683 |
| F4 48 | 4 | 6083.262136 | 0.04447478 | 1.471220099 | 3 | 3.42 | 0.0035 | 0.961484397 | 1.414555 |
| F4 53 | 4 | 4677.228272 | 0.066146604 | 1.131174703 | 3 | 3.43 | 0.0061 | 0.964295755 | 1.090787 |
| F5 38 | 4 | 2214.953776 | 0.072334608 | 0.535680436 | 3 | 3.605 | 0.0025 | 1.013494518 | 0.542909 |
| F5 43 | 4 | 2718.777308 | 0.057053435 | 0.657528762 | 3 | 3.668 | 0.0044 | 1.031206073 | 0.678048 |
| F5 48 | 4 | 3930.042162 | 0.049012705 | 0.950469812 | 3 | 3.648 | 0.0132 | 1.025583357 | 0.974786 |
| F5 53 | 4 | 3813.991401 | 0.053696917 | 0.922403257 | 3 | 4.016 | 0.0042 | 1.129041327 | 1.041431 |

**Table S2.**
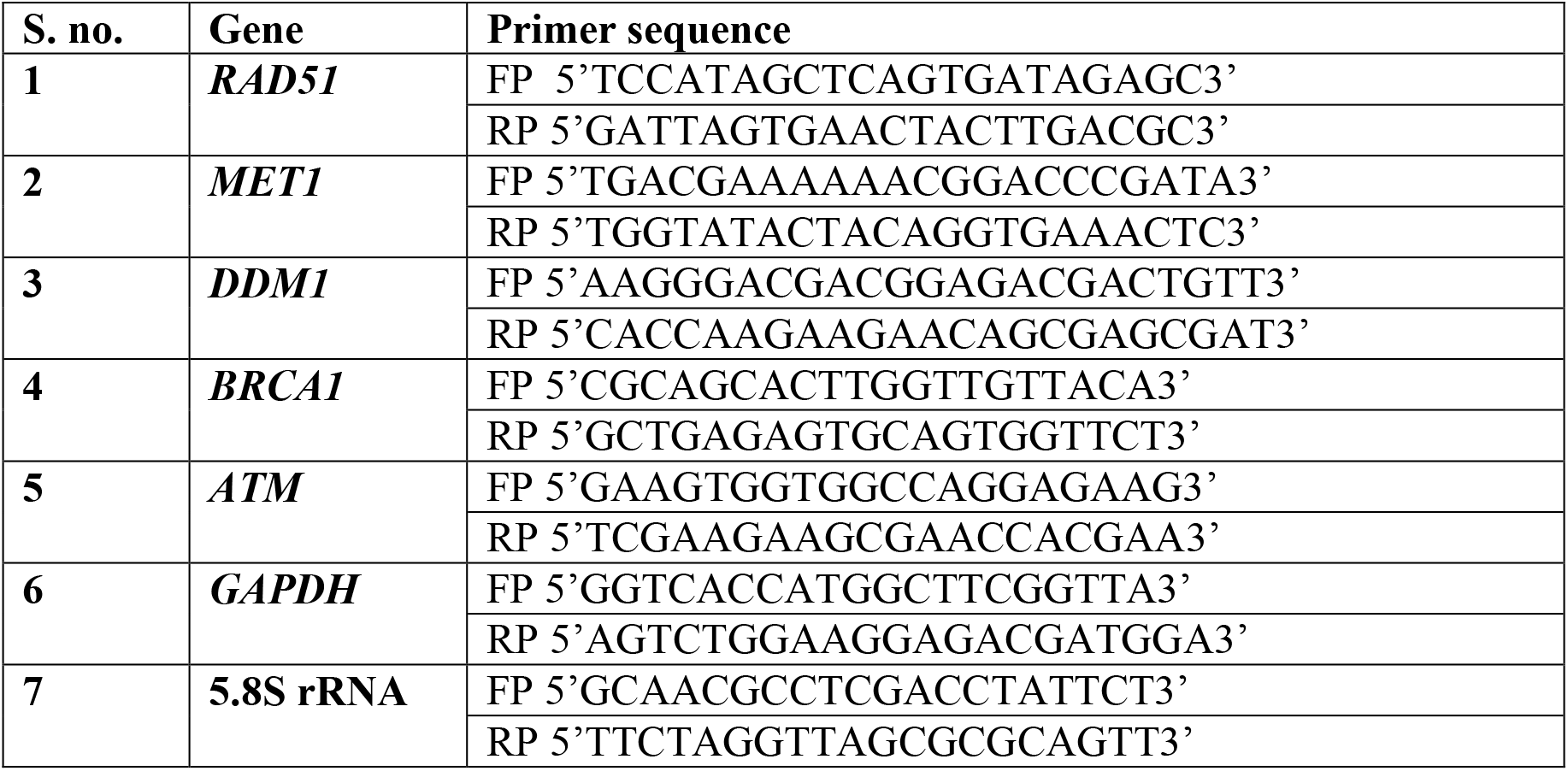
List of primers used for realtime PCR and their sequence

