## Supplementary figures and images for "Persistence of parental age effect on somatic mutation rates across generations in *Arabidopsis*"

### Supplemental table 1

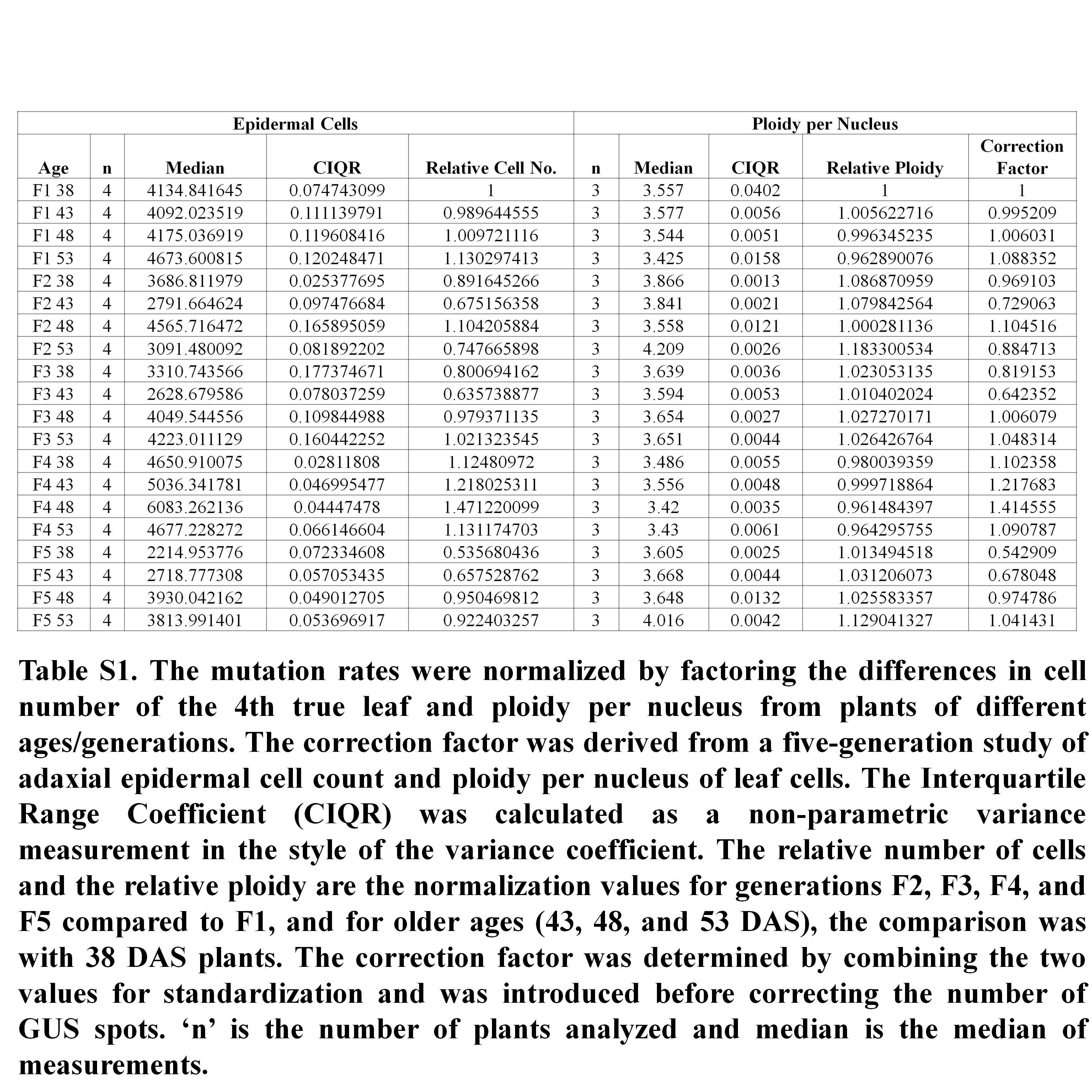

### Supplemental table 2

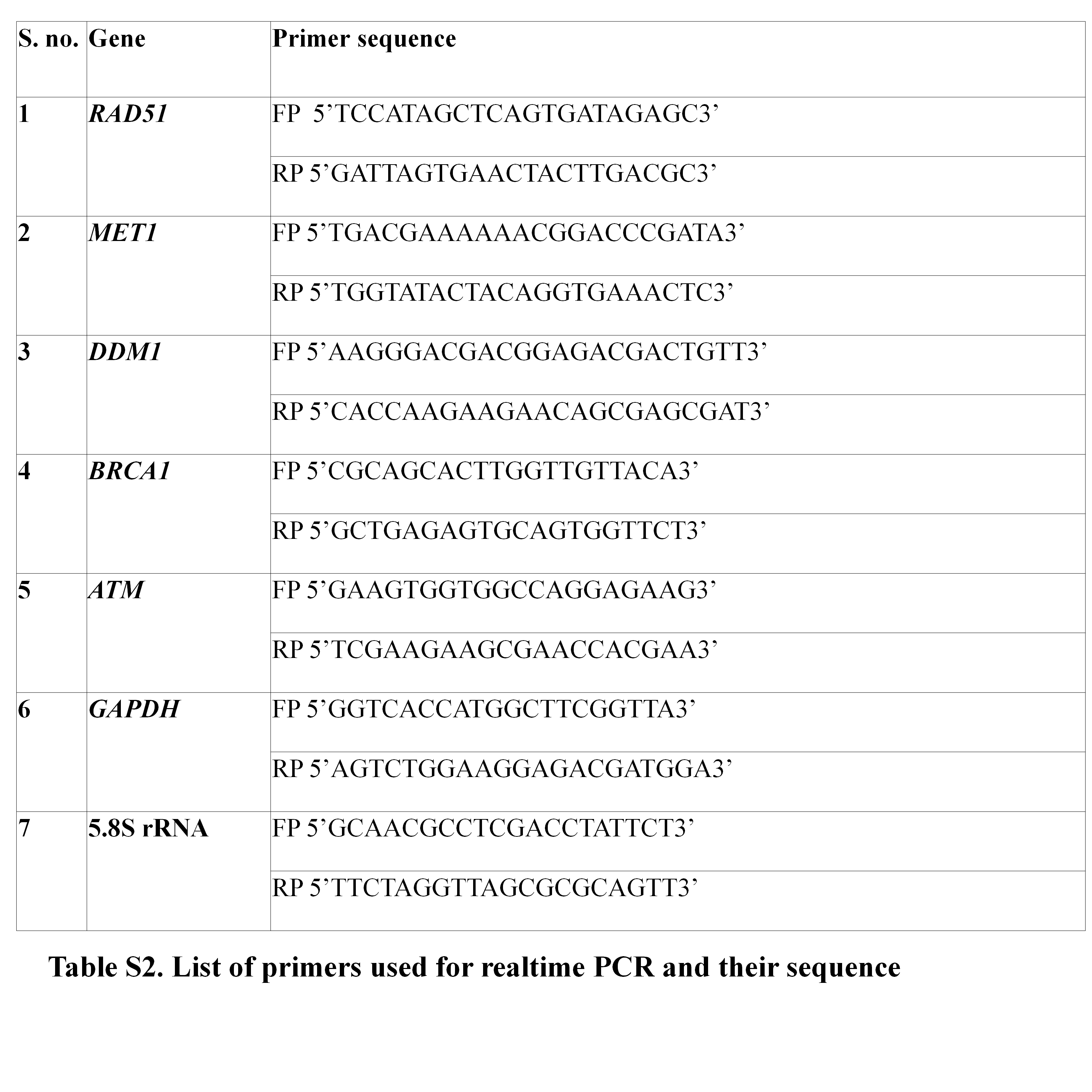
